# Copy number flexibility facilitates heteroresistance to increasing antibiotic pressure and threatens the beta-lactam pipeline

**DOI:** 10.1101/2022.06.07.494873

**Authors:** Jacob E. Choby, Tugba Ozturk, Carter N. Abbott, Christina Nnabuife, Jennifer M. Colquhoun, Sarah W. Satola, Philip N. Rather, Timothy Palzkill, David S. Weiss

## Abstract

It is unclear how bacteria rapidly adapt to resist novel drugs upon their introduction. Heteroresistance (HR) is a form of antibiotic resistance where a phenotypically unstable minority resistant subpopulation co-exists with a susceptible population. We observed HR to cefiderocol, a novel beta-lactam developed to resist beta-lactamases including extended-spectrum-beta-lactamases (ESBLs), among clinical isolates collected before its use. The resistant subpopulation in *Enterobacter* was a continuum; increasing gene copy number of an otherwise ineffective ESBL mediated increased resistance in decreasing numbers of cells. The factors that control the magnitude of amplification are unclear. We observed that ESBL activity controlled the level of amplification, thus increased copy number can compensate for poor enzymatic activity. A *Klebsiella* isolate from a clinical treatment failure also demonstrated amplification, highlighting the relevance of this phenomenon. These data provide insights into factors controlling dynamics of HR and how bacteria use phenotypic resistance to flexibly confront new antibiotic threats.

## INTRODUCTION

Antibiotic resistance is a growing crisis causing over 1 million worldwide deaths in 2019^1^. Without significant intervention, this number is predicted to rise to 10 million annual deaths by 2050^2^. The staggering death toll can be partly attributed to the widespread dissemination of antibiotic resistance, the rapid emergence of resistance to novel antibiotics, and a lagging pipeline for the development of novel classes of antibiotics. Variations of beta-lactams and beta-lactamase inhibitors have dominated the antibiotic pipeline in part because of their efficacy, low toxicity, and the many obstacles^3^ that hinder the development of completely novel antibiotic classes. In 2020, 47% of outpatient antibiotic prescriptions in the United States were beta-lactams, nearing 100 million^4^. Five of nine new antibiotics expected to treat highly resistant Gram-negative pathogens and approved from 2014 to 2020 were beta-lactams, and as of 2020, 7/7 antibiotics in Phase III testing were beta-lactams^5^. This reliance on beta-lactam antibiotics is a healthcare vulnerability since resistance emerges quickly following clinical introduction, for example, resistance to ceftazidime-avibactam was described less than a year following introduction^6^. However, the reasons for this rapid resistance are unclear^7^.

Cefiderocol is a recently FDA-approved beta-lactam consisting of a siderophore linked to a novel hybrid cephalosporin. The siderophore portion facilitates the transport of cefiderocol into the bacterial cell via siderophore receptors, while the late generation cephalosporin (a subclass of beta-lactams) moiety consisting of parts of ceftazidime and cefepime was designed to circumvent hydrolysis by beta-lactamases which cleave and inactivate beta-lactams. Enzymatic assays with cefiderocol showed relative resistance to hydrolysis by Ambler class A and D (serine) and B (metallo-) carbapenemases^8,9^, as well as class C AmpC-beta-lactamases^10^. Broad screens of clinical isolates using conventional antimicrobial susceptibility testing (broth microdilution; BMD) revealed low minimum inhibitory concentrations (MIC) against extended-spectrum beta-lactamase (ESBL)-, metallo-beta-lactamase-, and carbapenemase-producing isolates^11–15^, suggesting cefiderocol would be an efficacious new antibiotic for difficult to treat infections caused by Gram-negative pathogens. However, in a recent clinical trial (CREDIBLE-CR) for the treatment of serious infections caused by carbapenem-resistant isolates, higher than expected rates of cefiderocol treatment failure occurred^16^. In particular, cefiderocol was associated with increased all-cause mortality in patients with infections caused by carbapenem-resistant *Acinetobacter baumannii* (CRAB).

Antibiotic treatment failure may be caused by heteroresistance (HR), a form of antibiotic resistance in which a minority population of resistant cells co-exists with a majority susceptible population (Figure S1A). Treatment with a given antibiotic prevents the growth of the susceptible cells while the resistant subpopulation grows and dominates the population, distinguishing this form of resistance from persistence or tolerance. When the antibiotic is removed, the resistant subpopulation returns to its original homeostatic frequency (Figure S1B). We and others have observed HR to all classes of antibiotics tested^17,18^ and have shown that HR can cause treatment failure in murine models of infection^17,19,20^.

To investigate the discordance between the widespread susceptibility to cefiderocol in laboratory testing and the underwhelming patient outcomes in clinical testing, we curated a collection of isolates representative of those in the CREDIBLE-CR trial, from patients pre-dating the clinical introduction of cefiderocol. We observed a substantial frequency of HR especially among CRAB isolates (>50%), and the rate of HR for each species tested closely matched their all-cause mortality rate in the clinical trial^21,22^. These data suggest that undetected cefiderocol HR may have contributed to treatment failure in the Phase III testing of this antibiotic.

The molecular basis of cefiderocol HR is unknown. In this study, we investigated mechanisms of cefiderocol HR in Gram-negative clinical isolates and found that beta-lactamase gene amplification generates a continuum of subpopulations resistant to cefiderocol, where decreasing numbers of cells have increasing copy number and thus resistance. Genetic inhibition or use of beta-lactamase inhibitors (BLI) to reduce beta-lactamase activity revealed a functional flexibility whereby enhanced copy number of an otherwise ineffective beta-lactamase overcame inhibition. Further, exposure to sub-breakpoint concentrations of a cefiderocol/BLI combination led to increases in ESBL copy number sufficient to resist the combination. Therefore, amplification enables cellular subpopulations to employ otherwise ineffective ESBLs to resist cefiderocol, and additional copy number increases facilitate adaptation to increasing antibiotic stress such as from novel BLIs. These data lead to novel insights into the population dynamics of HR and how unstable, phenotypic resistance can be used by bacteria to flexibly confront new antibiotic threats, even in the absence of stable evolutionary changes. This phenotypic plasticity is undetected during antibiotic development and is a threat to the antibiotic pipeline, which is dominated by beta-lactams/BLIs.

## RESULTS

### Increased ESBL copy number generates cefiderocol heteroresistance

Using population analysis profile (PAP; Figure S1C-D) to quantify the frequency of resistant cells by plating dilutions at an array of drug concentrations, we identified cefiderocol HR in *Enterobacter cloacae* complex strain RS isolated from a renal transplant patient^20^. The frequency of the RS cefiderocol resistant subpopulation at the clinical breakpoint concentration (which differentiates susceptible and resistant isolates^23^) was 1 in 20,000. In contrast, the entire population of cells of susceptible isolate ESBL89 was killed at concentrations below the breakpoint (Figure 1A).

**Figure 1.**
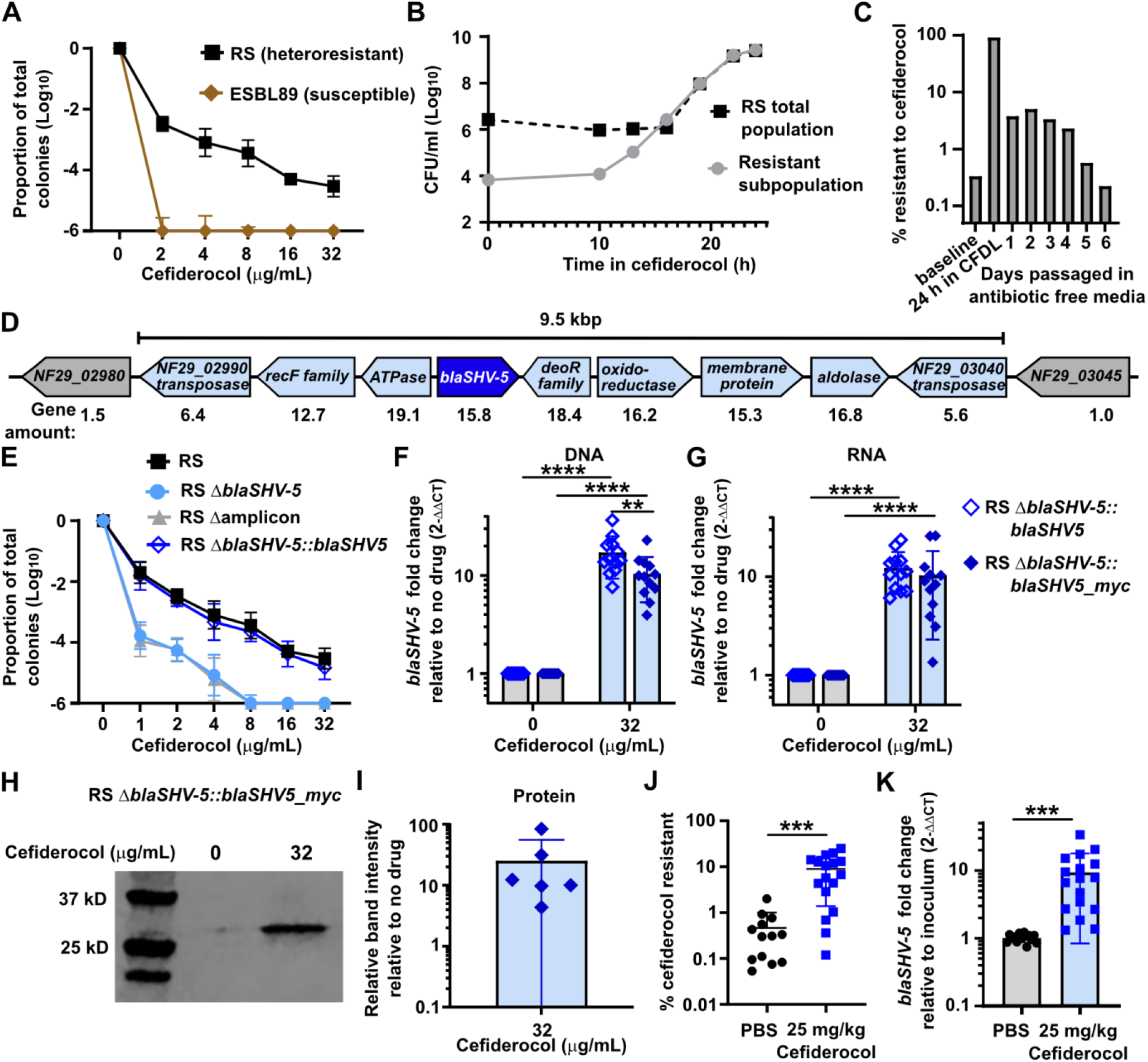
Increased *blaSHV-5* copy number generates cefiderocol heteroresistance in Enterobacter strain RS. (**A**) Population analysis profile (PAP) of strains RS and *Klebsiella pneumoniae* ESBL89 plated on Mueller-Hinton agar (MHA) containing cefiderocol; the proportion of surviving colonies is quantified relative to MHA containing 0 cefiderocol, the mean and standard deviation are shown from two independent experiments with 3 biological replicates (RS) or 1 biological replicate (ESBL89) each shown. (**B**) Time kill of strain RS: growth in media containing 16 μg/mL cefiderocol over time and quantification of surviving colony forming units on MHA (RS total population) and MHA containing cefiderocol (resistant subpopulation) from a single representative replicate. (**C**) Quantification of the cefiderocol resistant subpopulation of strain RS after growth in media alone (baseline), subcultured into cefiderocol and grown for 24 h, and subcultured every 24 h into fresh media without cefiderocol. At the end of each growth, an aliquot was diluted and plated onto MHA containing cefiderocol to quantify the resistant subpopulation, from a single representative replicate. (**D**) The region of the RS chromosome amplified in cells surviving growth in cefiderocol is shown, with gene abundance shown below. (**E**) PAP on cefiderocol of RS, the Δ*blaSHV-5* and Δamplicon (Δ*NF29_02990-NF29_03045*) isogenic mutants, and the Δ*blaSHV-5* mutant complemented with *blaSHV-5* at the native site; the mean and standard deviation are shown from two independent experiments with 3 biological replicates each. (**F-G**) *blaSHV-5* gene abundance quantified from DNA (**F**) or *blaSHV-5* transcript abundance quantified from RNA (**G**) by qPCR from colonies collected from MHA containing 32 μg/mL cefiderocol normalized to no cefiderocol for each replicate from the strains indicated. Mean and standard deviation are shown, from two experiments with 6 biological replicates each. ** indicates p<0.01, **** indicates p<0.0001 by one-way ANOVA, [ F (3, 44) = 34.17 for (F), F (3, 44) = 18.29 for (G)] with Sidak’s multiple comparisons test. (**H**) Representative anti-myc immunoblot from total cellular lysate from colonies collected from MHA containing 0 or 32 μg/mL cefiderocol of strain RS Δ*blaSHV-5*::*blaSHV-5_myc*. SHV-5_myc is 30.09 kD following cleavage of signal sequence. **I**) anti-myc immunoblot band density analysis with LiCor analysis, showing the relative ratio of the band intensity from samples grown on 32 μg/mL cefiderocol relative to no drug, from six biological replicates across two independent experiments. Full blots and additional images corresponding to H-I are shown in Supplemental Figure 5. (**I-J**) Outcomes of infection of *Galleria* larvae and cefiderocol treatment, (**I**) The frequency of the subpopulation resistant enumerated on agar containing 2 μg/mL cefiderocol with mean and standard deviation shown. (**J**) *blaSHV-5* gene abundance from CFU collected from agar containing 2 μg/mL cefiderocol, normalized to infection inoculum, mean and standard deviation are shown. For I-J, each symbol indicates a single larva, from two independent experiments, n=13 for PBS treated and n=17 for cefiderocol treated; *** indicates p<0.001 by unpaired t-test with Welch’s correction [t=4.610, df=16.21 for (J) and t=4.039, df=16.01 for (K)].

We investigated the dynamics of the RS resistant subpopulation during cefiderocol treatment. The resistant subpopulation replicated in cefiderocol, demonstrating that it is not a population of persister cells, which do not replicate in the presence of antibiotic^24^ (Figure 1B). A second hallmark of HR is phenotypic instability of the resistant subpopulation. To test whether the resistant subpopulation was stable or unstable, we grew RS in cefiderocol, and then serially passaged in the absence of the drug. We observed enrichment of the resistant subpopulation in cefiderocol and the subsequent return to the pre-selection frequency in its absence, consistent with unstable heteroresistance^18^ (Figure 1C). Together, these data indicate that RS exhibits HR to cefiderocol.

We next set out to determine if the resistant subpopulation was genetically distinct from the majority susceptible population. We sequenced the genome of RS after growth in broth with 32 µg/mL (two-times the breakpoint) cefiderocol (>99% resistant cells) or no drug (>99% susceptible cells). Bioinformatic analysis revealed a 9.5 kilobase pair (kbp) region in the chromosome of the resistant cells grown in cefiderocol that was present at an increased frequency of ∼15x (which we describe as having an increased copy number or being “amplified”, i.e. exhibiting gene amplification). This region included *blaSHV-5*, encoding a class A extended-spectrum beta-lactamase (ESBL; Figure 1D). We collected colonies of the resistant subpopulation from agar containing cefiderocol, and used qPCR to confirm that several genes within the amplified region had elevated copy number, but no change was detected in copy number of genes flanking the amplified region (Figure S2). We also selected 15 resistant colonies from agar plates containing cefiderocol, grew each in broth without the drug, and then performed qPCR to quantify the number of *blaSHV-5* copies. The extent of *blaSHV-5* gene amplification in the resistant colonies was always greater than 1 but with variable magnitude, consistent with *blaSHV-5* copy number heterogeneity in the population (Figure S3). Next, we hypothesized that the increased copy number is the result of tandem gene amplification, and exists at a low frequency prior to antibiotic exposure. Tandem duplication and amplification by recombination during DNA replication^25^ may occur between the identical co-directional *NF29_02990* and *NF29_03040* genes flanking the *blaSHV-5* amplicon. After growth of RS in cefiderocol, we observed a PCR product consistent with the duplication event (Figure S4). Importantly, evidence for the tandem duplication was detected in RS prior to cefiderocol exposure, albeit at a greatly reduced intensity, consistent with a pre-existing, minor subpopulation of cells exhibiting *blaSHV-5* gene amplification. These data identify increased copy number of a region including a beta-lactamase within the resistant subpopulation of cells in a cefiderocol HR clinical isolate.

To test whether this amplified region was responsible for cefiderocol HR, we generated in-frame unmarked deletions of the entire amplicon region (Δamplicon) or *blaSHV-5* alone (Δ*blaSHV-5*) and tested the strains by PAP. The resistant subpopulation was not present in the Δamplicon mutant nor in the Δ*blaSHV-5* mutant (Figure 1E). Complementation by reintroduction of *blaSHV-5* at its native site in the Δ*blaSHV-5* mutant restored the resistant subpopulation (Figure 1E). Thus, *blaSHV-5* is required for the presence of the cefiderocol resistant subpopulation.

Having established that the resistant subpopulation has elevated *blaSHV-5* copy number, we sought to test whether this increased gene copy resulted in greater SHV-5 enzyme abundance. The native *blaSHV-5* was replaced with a version encoding a C-terminal myc epitope, which retains wild-type function (Figure S5A). Resistant colonies from strains encoding the native or myc-tagged SHV-5 were collected from agar. The resistant colonies growing on 32 ug/ml cefiderocol demonstrated enhanced *blaSHV-5* DNA copy number (Figure 1F) as well as increased mRNA transcript abundance relative to colonies grown on agar without drug (Figure 1G). Under these same conditions, a robust increase in SHV-5 enzyme was apparent in the resistant population selected by growth on cefiderocol (Figure 1H-I). These data establish that the resistant subpopulation, which carries more copies of the *blaSHV-5* gene, produces more SHV-5 enzyme than the predominantly susceptible population in the absence of cefiderocol. Furthermore, after growth in cefiderocol, the *blaSHV-5* mRNA and protein levels in the resistant subpopulation correlated well with the magnitude of gene amplification (∼10-20x increase compared to cells grown without drug). We established a model of *Enterobacter* infection in *Galleria* waxworm moth larvae^26^ to test whether *in vivo* cefiderocol treatment selected for the resistant subpopulation with elevated *blaSHV-5* gene copy. Following infection and cefiderocol treatment, we observed an approximately 20-fold increase in the frequency of the resistant subpopulation (Figure 1J). We then quantified *blaSHV-5* copy number in the resistant subpopulation and observed a 10-fold increase following *in vivo* cefiderocol treatment (Figure 1K). Thus, we have observed *in vivo* selection of the cefiderocol resistant subpopulation with a corresponding increase in *blaSHV-5* gene frequency. Together, these data establish beta-lactamase gene amplification as being responsible for a dynamic resistant subpopulation which is selected by antibiotic treatment.

### Cefiderocol heteroresistance comprised of a continuum of resistant subpopulations

These data led to a model where in the absence of cefiderocol, the majority of cells have a single copy of the ESBL encoding amplicon and a minority exhibit *blaSHV-5* gene amplification. In contrast, in the presence of cefiderocol there is enrichment of the cells with *blaSHV-5* gene amplification. To further test this model, we grew RS with or without increasing concentrations of cefiderocol and quantified *blaSHV-5* gene frequency. Interestingly, as the concentration of cefiderocol increased and the frequency of the resistant subpopulation decreased, the average number of *blaSHV-5* copies in the surviving resistant subpopulation increased (Figure 2A). At 8 µg/mL, 1 in 5,000 cells survived with an average gene frequency of 2, whereas at 32 µg/mL, 1 in 150,000 cells survived with an average gene frequency of 20 (Figure 2B). This suggested that the cells surviving in increasing concentrations of cefiderocol require increasing *blaSHV-5* amplification to withstand the additional stress. Furthermore, these data surprisingly indicate that the cefiderocol resistant subpopulation is actually a continuum of subpopulations with increasing *blaSHV-5* gene frequency and resistance levels. This further highlights that among the resistant cells surviving at a given concentration of cefiderocol, the level of *blaSHV-5* amplification is not uniform. For example, at 8 µg/ml cefiderocol, a majority of surviving cells would have an average *blaSHV-5* frequency of 2 (as mentioned above), but minorities of surviving cells would have average *blaSHV-5* frequencies of between 6 and 20 and be capable of also surviving at 16 µg/ml or 32 µg/ml, respectively (Figure 2C).

**Figure 2.**
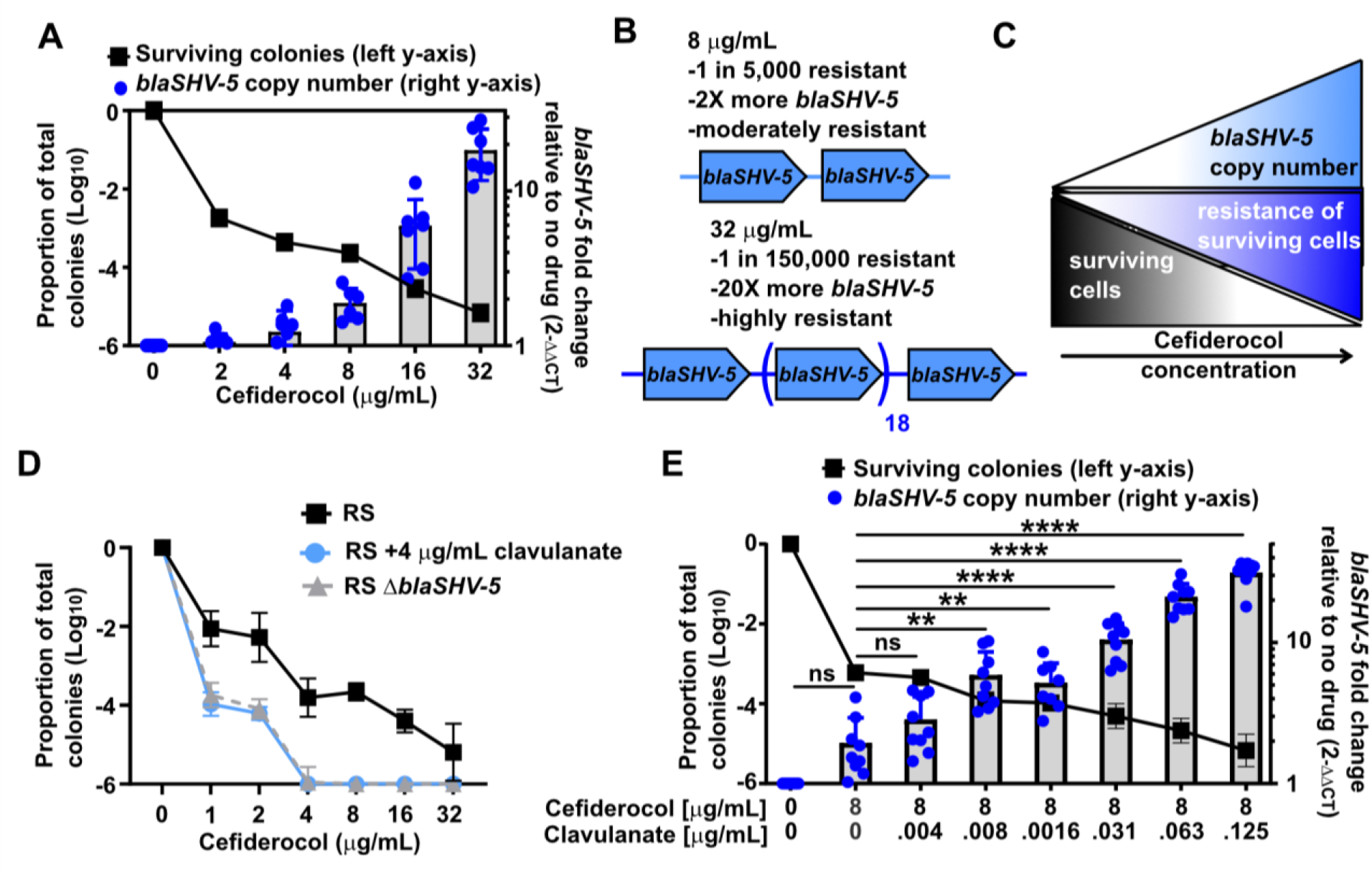
Amplification generates a subpopulation continuum responsive to inhibition. (**A**) PAP of RS plated on cefiderocol, with the proportion of surviving colonies in the black line on the left y-axis, with mean and standard deviation shown. *blaSHV-5* abundance was quantified from the samples collected at each concentration. Each dot indicates a biological replicate, and bars show the mean and standard deviation graphed on the right y-axis, from two independent experiments with 3-4 biological replicates each. * indicates p<0.05, ** p<0.01 by mixed-effects ANOVA analysis with Geisser-Greenhouse correction and Dunnet’s multiple comparison test. (**B**) Details of the populations collected on 8 µg/mL or 32 µg/mL cefiderocol are shown. (**C**) A model of the RS population: as the concentration of cefiderocol increases, the proportion of the surviving cells decreases. The cefiderocol resistance of the surviving resistant cells increases with an increase in the *blaSHV-5* abundance. (**D**) PAP of strain RS or the isogenic Δ*blaSHV-5* mutant plated on cefiderocol or cefiderocol and 4 µg/mL clavulanate. The mean and standard deviation is shown from two independent experiments with 3 biological replicates each. (**E**) Strain RS was plated on MHA containing cefiderocol and clavulanate as indicated. The proportion of the surviving population is the line graph with the left y-axis with means and standard deviation. The corresponding gene abundance is graphed on the right y-axis, where each point indicates a biological replicate from two independent experiments with 3 or 6 biological replicates each (n=9 total) and bars indicate mean and standard deviation. ** indicates p<0.01, **** p<0.0001 by mixed-effects ANOVA, F (1.909, 14.72) = 112.2 with Geisser-Greenhouse correction and Dunnet’s multiple comparison test.

The fact that the magnitude of gene amplification in the resistant subpopulation correlates with cefiderocol concentration (Figure 2A) suggested that strains require more beta-lactamase function to resist higher concentrations of cefiderocol. Since beta-lactamase inhibitors (BLIs) reduce the activity of beta-lactamases, we hypothesized that if a strain were to survive in a given concentration of cefiderocol plus BLI, it would require more beta-lactamase function to survive than in the same concentration of cefiderocol alone. To test this hypothesis, we used the BLI clavulanate which eliminates the highly resistant subpopulation when combined with cefiderocol (Figure 2D). We therefore selected resistant colonies of RS on 8 µg/mL cefiderocol and increasing concentrations of clavulanate (Figure 2E). As the frequency of the resistant subpopulation decreased with increasing clavulanate, we observed a concomitant increase in *blaSHV-5* amplification (Figure 2E). To quantify the magnitude of *blaSHV-5* amplification in cells grown on cefiderocol/clavulanate by an alternative assay, we performed droplet digital PCR and observed *blaSHV-5* amplification to a similar extent as qPCR (Figure S6). These results, therefore, links *blaSHV-5* copy number and the extent of SHV-5 activity, because when more clavulanate is present, the enzyme will be inhibited to a greater degree and more copies are required to survive. These data suggest that as the overall antibiotic stress increases, this strain can survive due to the flexibility afforded by gene amplification.

### Inhibition of beta-lactamase activity leads to a compensatory increase in ESBL copy number

Our observation that the addition of a BLI led to increased ESBL gene amplification (Figure 2E) is consistent with our hypothesis that in order to survive a given cefiderocol exposure, cells would require more copies of a beta-lactamase if it has lower activity. To test this hypothesis, we evaluated the impact of a point mutation in SHV-5 (M69I) which was previously demonstrated to reduce activity towards the cephalosporins cefotaxime and cephalothin^27^. We expressed and purified the wild-type and M69I point mutant SHV-5 enzymes from *E. coli*. Steady-state kinetic parameters indicated that SHV-5 hydrolyzes cefotaxime and cephalothin with *k*_cat_/*K*_M_ values of 8 x 10^5^ and 1 x 10^6^ M^-1^s^-1^ (Figure S7). Further, SHV-5 was found to hydrolyze cefiderocol with a *k*_cat_/*K*_M_ of 9 x 10^3^ M^-1^s^-1^, which is greatly reduced compared to cefotaxime and cephalothin, but clearly indicates a low level of active hydrolysis (Figure S7). In contrast, SHV-5 M69I exhibited *k*_cat_/*K*_M_ values of 2.3 x 10^4^ and 8 x 10^4^ M^-1^s^-1^ for cefotaxime and cephalothin, respectively. We were unable to accurately determine *k*_cat_ and *K*_M_ for catalysis with cefiderocol due to a high *K*_M_ but were able to determine *k*_cat_/*K*_M_ as 1.2 x 10^3^ M^-1^s^-1^. Therefore, the M69I substitution reduced catalytic efficiency by 35- and 10-fold for cefotaxime and cephalothin, respectively, and 7-fold for cefiderocol.

We also generated an M69I point mutant version of *blaSHV-5* which was inserted at the Δ*blaSHV-5* locus, creating Δ*blaSHV-5*::*blaSHV-5* M69I (*blaSHV-5* M69I, encoding SHV-5 M69I). Δ*blaSHV-5*::*blaSHV-5* (*blaSHV-5*) was used as a control strain expressing parental, wild-type *blaSHV-5*. We performed PAP and observed that the strain encoding SHV-5 M69I harbored subpopulations with reduced resistance (only surviving up to 4 µg/ml cefiderocol) as compared to the strain encoding parental SHV-5 (which survives up to at least 32 µg/mL cefiderocol) (Figure 3A). However, the strain encoding SHV-5 M69I exhibited greater resistance than the Δ*blaSHV-5* mutant which only survived up to 2 μg/mL (Figure 3A). We next quantified the average level of *blaSHV-5* amplification in bacteria expressing parental or M69I SHV-5 grown with or without 4 µg/mL cefiderocol, a concentration at which at least some of the cells of each strain could survive (Figure 3A). We observed a trend towards increased amplification of the gene encoding SHV-5 M69I compared to wild-type SHV-5 (Figure 3B).

**Figure 3.**
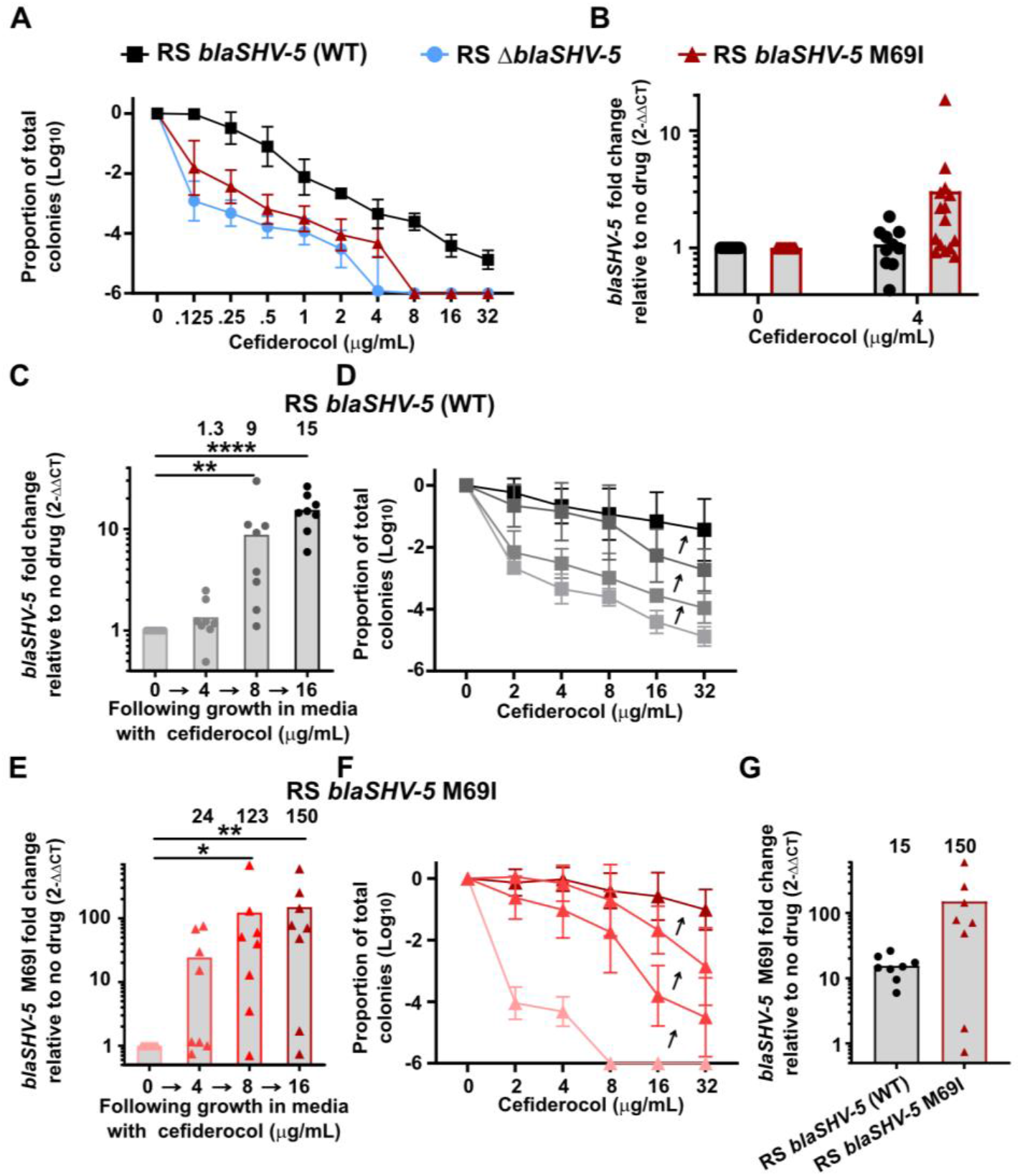
Inhibition of beta-lactamase activity leads to a compensatory increase in *blaSHV-5* copy number. (**A**) PAP of *Enterobacter* RS strains plated on cefiderocol. Genotypes are Δ*blaSHV-5*, Δ*blaSHV-5*::*blaSHV-5* (referred to as *blaSHV-5* (WT)), and Δ*blaSHV-5*::*blaSHV-5 M69I* (*blaSHV-5* M69I). Shown are the means and standard deviation of three independent experiments with 3 biological replicates each. (**B**) *blaSHV-5* and *blaSHV-5 M69I* amplification from surviving colonies on 4 µg/mL cefiderocol in (**A**), each point indicates a replicate from three independent experiments, 10-15 replicates total for each strain.**** indicates p<0.0001 by two-way ANOVA with Sidak’s correction for multiple comparisons. (**C**) *blaSHV-5* abundance after resistant colonies were collected on MHA containing cefiderocol and grown in broth containing the same concentration indicated. (**D**) PAP of samples from panel (**C**). (**E**) *blaSHV-5* M69I abundance after resistant colonies were collected on MHA containing cefiderocol and grown in broth containing the same concentration indicated. (**F**) PAP of samples from panel (**E**). (**G**) *blaSHV-5* M69I fold change following growth in 16 µg/ml cefiderocol, from panels C and E, highlighting the difference in magnitude of amplification. **C-G** show the mean from 3 independent experiments with 2-3 biological replicates each, with each dot indicating a biological replicate in C, E, and G and standard deviation shown in D, and F. For C, and E, * indicates p<0.05, ** p<0.01, *** p<0.001, and **** p<0.0001 by Kruskal-Wallis test (used because samples failed D’Agostino & Pearson test for normality) with Dunnet’s correction for multiple comparisons.

Taken together, we have established that 1] there is a pre-existing continuum of subpopulations with differing levels of *blaSHV-5* amplification and corresponding cefiderocol resistance levels present in RS (Figure 2A) and 2] that the magnitude of amplification inversely correlates with the extent of SHV-5 function, which can be inhibited through chemical or genetic means (Figures 2E, 3B). These data clearly indicate that the extent of *blaSHV-5* amplification in RS is flexible and not fixed. Therefore, we set out to test whether serial passage of the cefiderocol resistant population could select for cells with increasing *blaSHV-5* copy number and corresponding resistance. We collected the resistant subpopulation of RS encoding wild-type SHV-5 from plates with 4 μg/mL cefiderocol (Figure 3B) and passaged sequentially to 8 µg/ml and 16 µg/mL cefiderocol, performing PAP and quantifying *blaSHV-5* gene frequency after each passage. Serial selection resulted in subpopulations with amplification levels that increased from an average of 1.3 to 15 (Figure 3C). The frequency of the resistant subpopulations in RS encoding wild-type SHV-5 increased in abundance with each passage in increasing concentrations of cefiderocol, yielding an approximate 3,000-fold increase in the frequency of the subpopulation resistant to 32 µg/mL after the final passage relative to the baseline (Figure 3D).

We next performed the same experiment with RS encoding SHV-5 M69I (*blaSHV-5* M69I). We observed a similar increase in *blaSHV-5* (M69I) gene amplification from baseline following passage (Figure 3E) and increases in the both the frequency of the resistant subpopulation and the highest concentration of cefiderocol on which the subpopulation survived (Figure 3F). The abundance of the gene encoding SHV-5 M69I was greater than that of SHV-5 at each passage step, and of note, the extent of gene amplification in the colonies following growth in 16 µg/mL cefiderocol was ten-fold greater for *blaSHV-5* M69I compared to *blaSHV-5* (Figure 3G). These data demonstrate a new paradigm in heteroresistance and copy number variation, where the level of activity of an amplified gene is a critical determining factor for the extent of amplification.

We further confirmed this finding by similarly generating Δ*blaSHV-5*::*blaSHV-5* S238G;K240E, converting SHV-5 to the parental, non-extended-spectrum beta-lactamase SHV-1 which has reduced activity towards cephalothin, ceftazidime, and cefotaxime relative to SHV-5^2^. This strain, *blaSHV-5* S238G;K240E (SHV-1) also demonstrated increased magnitude of amplification at every serial exposure to cefiderocol, which resulted in the rare subpopulation (∼1 in 100,000) resistant to only 4 µg/ml cefiderocol at baseline, shifting to a high proportion of cells (∼1 in 10) resistant to 32 µg/mL cefiderocol after the final exposure (Figure S8). These data together confirm that the beta-lactamase mutant strains with reduced activity require enhanced *blaSHV* gene amplification to survive on cefiderocol, and underscore the potential for vast *blaSHV-5* amplification and resulting increases in cefiderocol resistance.

In order to test the applicability of these findings beyond *Enterobacter*, we identified cefiderocol HR isolates of carbapenem-resistant *Acinetobacter baumannii* exhibiting beta-lactamase amplification and tested whether the extent of gene amplification increased upon inhibition of the beta-lactamase with a BLI. Using two representative isolates, we observed that the extent of amplification of the beta-lactamase gene was increased following growth in cefiderocol with the beta-lactamase inhibitor avibactam relative to growth in cefiderocol alone (see Extended Data, Figure S9-11). Thus, the link between beta-lactamase activity and the extent of gene amplification is shared in at least two antibiotic-resistant Gram-negative species.

### ESBL copy number increase in response to greater antibiotic stress overcomes beta-lactamase inhibitors

The observations that serial exposure to cefiderocol selected for an increase in *blaSHV-5* gene amplification and greatly increased the frequency and resistance level of the resistant subpopulations led us to hypothesize that similar exposure might drive sufficient amplification of *blaSHV-5* to overcome inhibition by clavulanate. We showed that the extent of *blaSHV-5* amplification in RS was greater with increasing concentrations of clavulanate (Figure 2E), but cells only survived up to 8 µg/mL cefiderocol/0.125 µg/mL clavulanate. We sequentially harvested resistant cells from the highest concentration of cefiderocol/clavulanate on which they survived, and subsequently grew them on higher concentrations. Repeating this process, we observed sequential survival on 8 µg/mL/0.06 µg/mL, 16 µg/mL/0.25 µg/mL, and 8µg/mL/4µg/mL cefiderocol/clavulanate (Figure 4A). When we quantified the extent of amplification of *blaSHV-5* in cells surviving in each of these three conditions, we detected corresponding increases from 21 to 40 to 63 copies (Figure 4A). After growth in each sequential condition, PAP was performed and we observed a stepwise increase in the frequency of the resistant subpopulation (Figure 4B). This demonstrates a proof of principle that amplification of a beta-lactamase can allow a strain to overcome escalation of beta-lactam therapy when an otherwise effective BLI is combined with a beta-lactam. We additionally observed that a single exposure of *A. baumannii* strain Mu1956 to cefiderocol/avibactam was sufficient to enrich for a subpopulation resistant to the clinical breakpoint concentration of each drug^23^ (Extended Data, Figure S12). These data highlight how rapidly an HR isolate can develop a subpopulation of cells resistant to cefiderocol plus an effective beta-lactamase inhibitor and how beta-lactamase gene amplification threatens both current beta-lactam/BLIs as well as future agents currently in the development pipeline.

**Figure 4.**
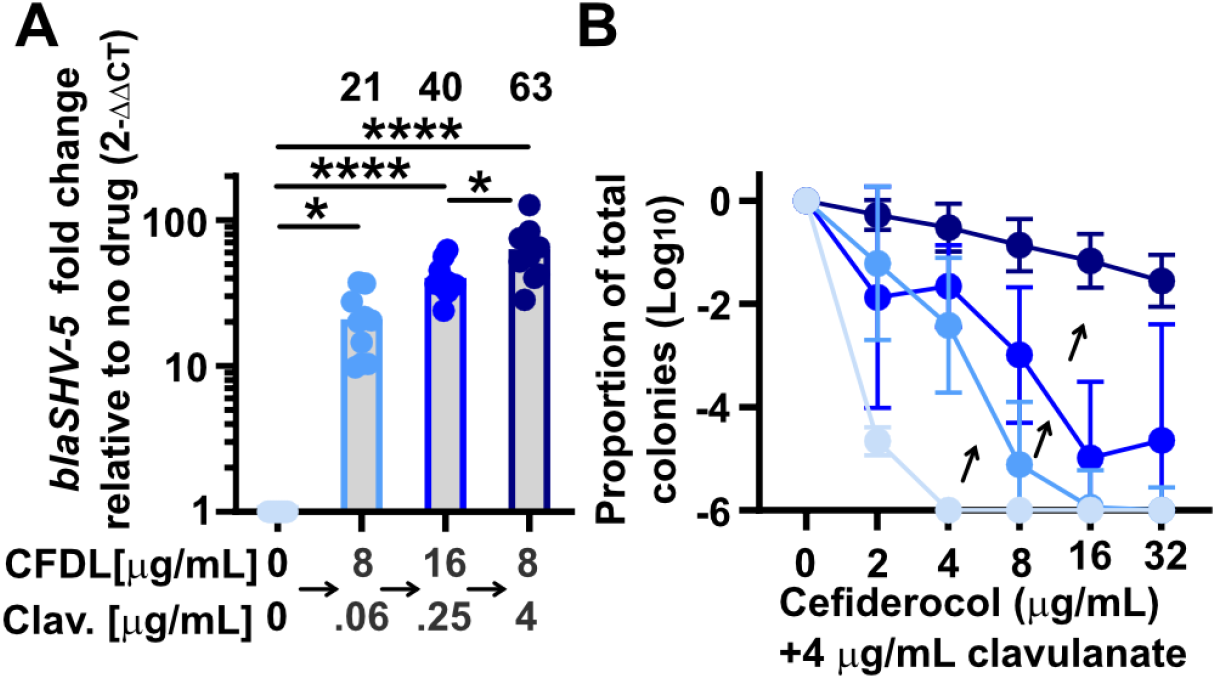
Cefiderocol/beta-lactamase inhibitor exposure increases amplification and results in a resistant subpopulation. **(A)** *blaSHV-5* abundance after serial exposure to cefiderocol/clavulanate. RS was plated on MHA alone or with 8 µg/mL cefiderocol and 0.06 µg/mL clavulanate and surviving colonies were collected and *blaSHV-5* abundance was quantified. A portion was subcultured into media with 8 µg/mL cefiderocol and 0.06 µg/mL clavulanate and grown for 24 h. The culture was plated on MHA containing 16 µg/mL cefiderocol and 0.25 µg/mL clavulanate. After 24 h, the resistant colonies were collected and *blaSHV-5* abundance was quantified. A portion was subcultured into media with 16 µg/mL cefiderocol and 0.25 µg/mL clavulanate and grown for 24 h. The culture was plated on MHA containing 8 µg/mL cefiderocol and 4 µg/mL clavulanate. After 24 h, resistant colonies were collected and *blaSHV-5* abundance was quantified. (B) PAP of RS plated on cefiderocol and 4 µg/mL clavulanate after growth in broth at each concentration of cefiderocol and clavulanate described in A. A-B show the mean of results from two independent experiments with 5 biological replicates each, shown as individual dots in A, standard deviation is shown in B. For A, * indicates p<0.05 and **** p<0.0001 by one-way ANOVA, F (3, 36) = 27.97, with Sidak’s correction for multiple comparisons.

### Beta-lactamase amplification in an isolate causing clinical cefiderocol failure

We have demonstrated that beta-lactamase amplification is a common phenomenon leading to cefiderocol HR, that amplification occurs *in vivo* following cefiderocol treatment (Figure 1K), and that cefiderocol HR correlates with and may lead to treatment failure in clinical trials^21,22^. We recently described a metallo-beta-lactamase producing *K. pneumoniae* strain isolated from a patient who failed cefiderocol therapy^29^ (Figure 5A). Although classified susceptible to cefiderocol by clinical testing, our analyses revealed that this strain is cefiderocol HR (Figure 5B). We sequenced this strain and identified a contig containing the metallo-beta-lactamase *blaNDM-5* (Figure 5C). When this isolate was treated with cefiderocol, the resistant subpopulation was enriched and *blaNDM-5* copy number increased (Figure 5D). These data demonstrate that *blaNDM-5* amplification occurs in an HR isolate from a patient who failed cefiderocol treatment, highlighting the potential for beta-lactamase amplification to subvert cefiderocol therapy in humans.

**Figure 5.**
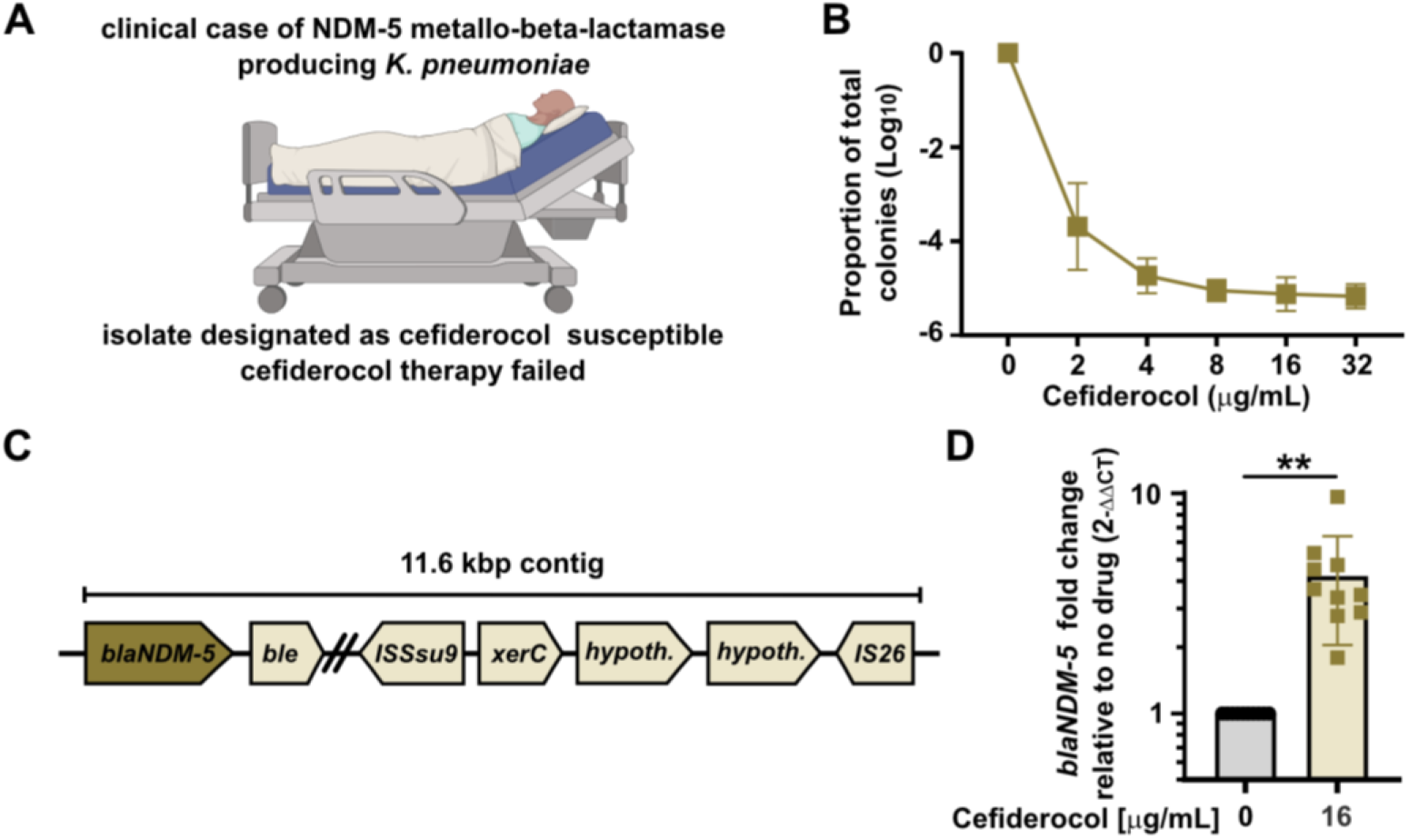
Beta-lactamase amplification in a heteroresistant isolate from a patient with failed cefiderocol therapy. (**A**) A patient was transferred to intensive care with septic shock and positive cultures with an NDM-5 metallo-beta-lactamase producing *K. pneumoniae*. Disk diffusion testing identified the isolate as cefiderocol susceptible and cefiderocol therapy was initiated, but failed. Subsequent testing identified cefiderocol heteroresistance. (**B**) Population analysis profile of the patient isolate on MHA containing cefiderocol, mean and standard deviation are shown from three independent experiments with 2-6 biological replicates each. (**C**) Depiction of the *blaNDM-5* encoding contig in the genome of the *K. pneumoniae* patient isolate. (**D**) *blaNDM-5* gene abundance following growth in broth containing 0 or 16 μg/mL cefiderocol with mean and standard deviation shown. Each symbol indicates a biological replicate from two independent experiments, with n=10 total for each group. ** indicates p<0.01 by unpaired t-test, t=4.687, df=9.000, with Welch’s correction.

## DISCUSSION

In this work, we describe a mechanism for cefiderocol HR, making novel and broad insights into the dynamics of HR mediated by resistance gene amplification and how this flexible phenotypic phenomenon can impact rapid adaptation to new beta-lactams/BLIs without the necessity for stable evolution. Despite being developed in part to resist ESBLs, we found that cefiderocol can be hydrolyzed by SHV-5 (Figure S7), and likely by other ESBLs when the genes are amplified to high copy number in subpopulations of resistant cells. In response to chemical inhibition by BLIs (Figure 2D-E) or genetic inhibition (Figure 3), additional increases in copy number of these otherwise ineffective ESBLs facilitated survival of HR isolates treated with cefiderocol. We observed that relative copy number in the resistant cells largely matched the relative mRNA abundance and enzyme abundance, directly connecting amplification to increased enzyme abundance (Figure 1F-I). Correspondingly, we observed an approximate 10-fold increase in copy number of the gene encoding SHV-5 M69I relative to wild-type SHV-5 following serial cefiderocol exposure (Figure 3G). This correlates in magnitude with the approximate 7-fold reduction in SHV-5 M69I catalytic efficiency relative to wild-type SHV-5 (Figure S7), suggesting that a proportional increase in beta-lactamase copy number and enzyme abundance can directly compensate for reduced enzymatic activity. These data demonstrate that the relative activity of a beta-lactamase is a critical determinant of its copy number during antibiotic treatment, greatly enhancing our fundamental understanding of copy number variation and heteroresistance.

A clinical isolate exhibiting HR to a given antibiotic is often described as harboring a subpopulation of resistant cells. Interestingly, studying cefiderocol HR we observed that this subpopulation can actually be a continuum of resistant subpopulations exhibiting a spectrum of beta-lactamase copy number and resulting cefiderocol resistance. Cells with modest increases in the number of copies of a given beta-lactamase are much more abundant than those with the greatest levels of gene amplification which are present at the lowest frequency. For example, in *Enterobacter* strain RS, 1 in 5,000 cells have an average of 2 copies of *blaSHV-5*, while 1 in 150,000 cells have an average of 20 copies (Figure 2B). The generation of this continuum of resistant cells is likely dependent on at least two factors; 1) baseline rate of duplication through homologous recombination by which gene amplification can occur and 2) constraint of gene amplification due to the increased fitness cost of carrying such a high level of a beta-lactamase encoding amplicon^30^. It is important to note that the fitness cost may only partly be due to the beta-lactamase itself but could also be due to other genes within the amplified region.

It is interesting to consider the dynamics of the continuum of resistant cells in HR isolates and what this means for the overall flexibility and fitness of such strains. During exposure to a given beta-lactam, the cells with the minimum level of gene amplification that is sufficient to facilitate survival would predominate. In this way, the cells with the lowest fitness cost possible are dominant. This strategy provides a strain with significant flexibility to balance survival and fitness. We expect this paradigm to be true for many of the multitude of genes capable of undergoing gene amplification across species and thus to be a foundational aspect of understanding HR. In addition to genes conferring resistance to beta-lactams, amplification of genes providing resistance to aminoglycosides^31,32^, sulfonamides^32^, tetracyclines^32^ and polymyxins^33^ have been identified in HR isolates.

Our findings of widespread cefiderocol heteroresistance paired with the apparent susceptibility of HR clinical isolates as classified by conventional antimicrobial susceptibility tests during the early testing of cefiderocol, suggest HR was overlooked during the development of this drug. One of the first assays in the development of a new beta-lactam is to determine if it withstands the activity of existing beta-lactamases, as assayed using *in vitro* biochemical tests of purified beta-lactamases. Significant data exists showing that cefiderocol resists the activity of most classes of beta-lactamases^8–10^. However, these assays using purified enzymes report enzymatic activity but do not take into account the possibility of beta-lactamase gene amplification and subsequent increases in enzyme abundance.

The next major assay used to evaluate the susceptibility of clinical isolates to a new beta-lactam is broth microdilution (BMD). In this assay, bacterial isolates are incubated with the beta-lactam in broth media and tested for growth to a visible optical density (cloudiness of the culture) by ∼16 hours. However, we observed that the isolates studied here are classified susceptible by BMD and their heteroresistance is not detected. If the isolates were first incubated with cefiderocol, however, which selects for the resistant subpopulation, the cefiderocol MIC by BMD was 16 or >64 µg/mL, which is considered resistant (Table S1). These data indicate that when the frequency of the resistant subpopulation is extremely low (in this case ∼1 in 10,000 cells), these resistant cells do not have sufficient time to grow to a visibly cloudy culture density in the 16 hour incubation time for BMD. Indeed, cefiderocol heteroresistance caused discrepancies in conventional susceptibility testing including BMD^34^. This highlights how HR can often be undetected by BMD and reveals an Achilles’ heel of the beta-lactam development/testing process.

This leads to a model of the novel beta-lactam development pipeline that is undermined at multiple steps by HR (Figure 6). By design, cefiderocol was resistant to *in vitro* hydrolysis by most beta-lactamases tested^8–10^. However, data presented in this manuscript demonstrate that increases in beta-lactamase copy number can generate a subpopulation of cells with enhanced resistance to cefiderocol (Figures 2 and 4). This population is present at low frequency and thus is not detected by BMD, the standard clinical susceptibility test for cefiderocol accepted by CLSI^35^. This led to Phase III testing for the treatment of infections caused by isolates that appeared to be susceptible to cefiderocol, but which resulted in relatively high rates of all-cause mortality. These high mortality rates correlated with the rates of cefiderocol HR in our clinical isolate collection^21,22^. During prolonged infection and treatment, we expect the resistant subpopulation to have sufficient time to grow and dominate the population, unlike the brief duration of growth before MIC determination by BMD. Our data demonstrating that a strain designated susceptible and associated with treatment failure in a human patient was cefiderocol HR (Figure 5) is consistent with this model. A cefiderocol heteroresistant isolate has also been recently described in a patient that underwent prolonged cefiderocol therapy, which ultimately failed^36^. Thus, we suggest that undetected HR resulting from beta-lactamase gene amplification may contribute to treatment failure of cefiderocol, as well as future antibiotics that rely on the beta-lactam moiety.

**Figure 6.**
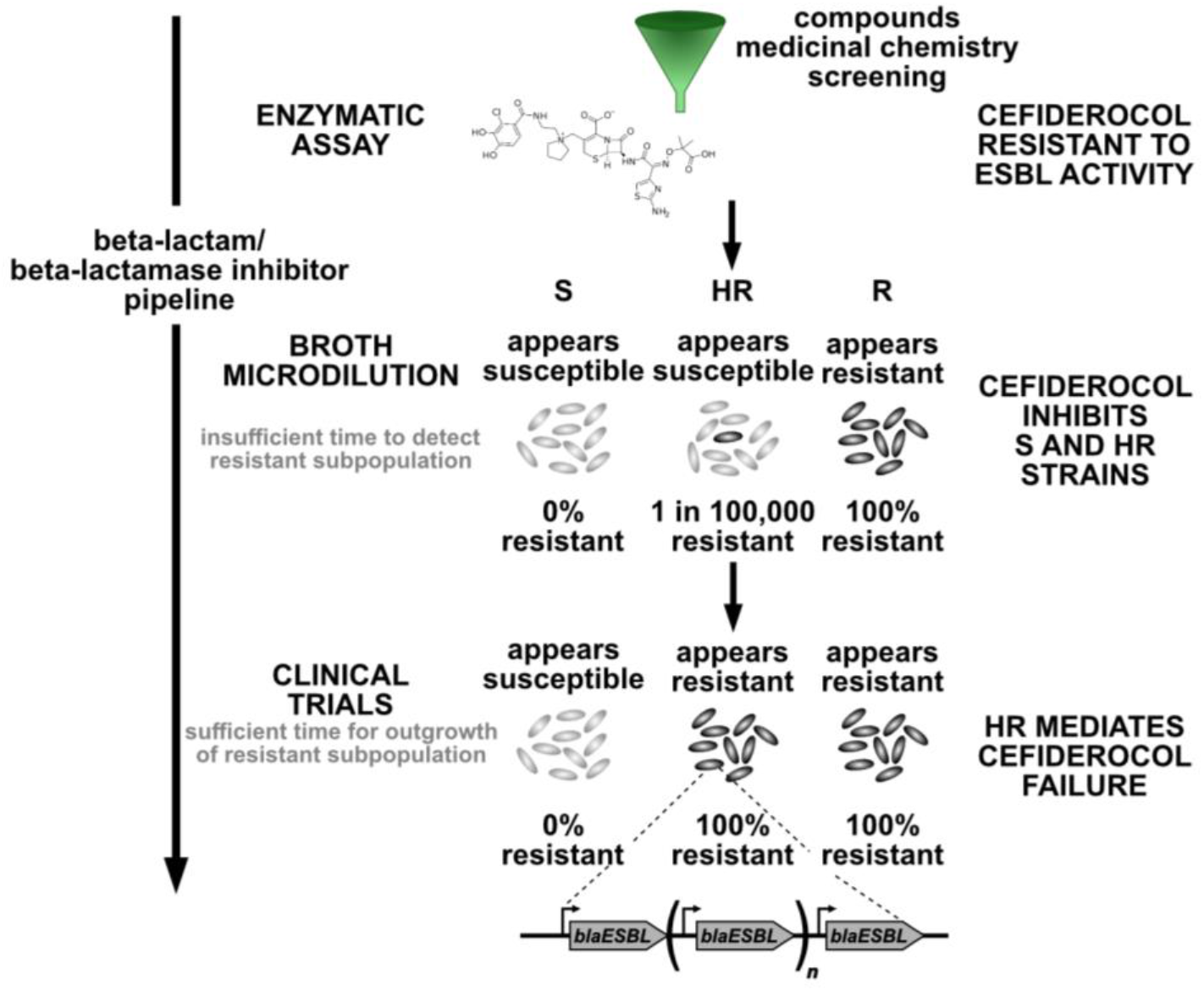
Heteroresistance threatens the beta-lactam/beta-lactamase inhibitor development pipeline. *In vitro* beta-lactamase enzymatic assays determined cefiderocol was resistant to extended-spectrum beta-lactamases (ESBL) and most carbapenemases tested. Broth microdilution (BMD) antimicrobial susceptibility testing assigned low cefiderocol minimum inhibitory concentrations (MIC) to both susceptible (S) and heteroresistant (HR) isolates, the latter of which harbored such a low frequency of resistant cells (e.g. 1 in 100,000) that they did not alter the overall MIC. BMD could detect only conventional resistance (R) in which 100% of the cells exhibit phenotypic resistance. HR is expected to cause treatment failure in patients because cefiderocol treatment selects for the resistant subpopulations which have amplifications of ESBL genes. These resistant cells become predominant during cefiderocol therapy and then can mediate treatment failure. In agreement with this model, unexpectedly high rates of treatment failure were observed in Phase III testing of cefiderocol, which correlated with rates of cefiderocol HR in surveillance studies. Taken together, the inability of the tests employed by the current antibiotic development pipeline (enzymatic assays and BMD) to detect HR leads drugs to which there are high rates of HR to progress to clinical testing where treatment failures may be observed. Incorporating testing for HR in the antibiotic development pipeline could potentially make the process more efficient and efficacious.

Beyond the potential to cause treatment failure of a beta-lactam to which a strain exhibits HR, we also investigated whether beta-lactamase gene amplification might contribute to adaptation to novel beta-lactams/BLIs to which a given HR isolate is initially susceptible. We observed that the BLIs clavulanate and avibactam could each render some HR isolates susceptible to cefiderocol (Figure S9). We subsequently demonstrated that just one exposure of such a strain to a ¼ breakpoint concentration of cefiderocol/avibactam could lead to enhanced beta-lactamase copy number, leading to resistance to the breakpoint concentrations (Figure S11). This shows how gene amplification can also mediate adaptation of a clinical isolate to a newly added BLI. Importantly, the clinical isolates studied here were collected before the clinical introduction of either cefiderocol or avibactam. Therefore, amplification of pre-existing beta-lactamases with suboptimal activity is a mechanism poised to mediate functional resistance and potential treatment failure of future beta-lactams/BLIs which have not yet even been introduced into the clinic. Importantly, this mechanism of phenotypic flexibility does not rely on new, stable evolution (e.g. of novel beta-lactamases with enhanced activity). These data highlight the paramount importance of screening for HR during the early stages of the drug development process, as well as during clinical susceptibility testing, to avoid potential treatment failures.

## METHODS

### Isolate information

*Enterobacter cloacae* complex strain RS is a clinical isolate previously described^20^.Carbapenem-resistant (CR) organisms were collected in Georgia, USA by the Georgia Emerging Infections Program as part of the CDC’s EIP Multi-site Gram-negative Surveillance Initiative (MuGSI): *Acinetobacter baumannii* (CRAB, from 2012-2015) and *Enterobacterales* spp. (CRE, from 2011-2015). Identification of cefiderocol heteroresistance in these isolates was previously described^21^. *A. baumannii* Mu1956 and Mu1984, along with the metallo-beta-lactamase producing *K. pneumoniae* genomes were assembled de novo and uploaded to NCBI Genbank (see Materials and Methods Table 1).

### Reagents

Mueller-Hinton agar (MHA; BD Difco), Mueller-Hinton broth (MHB; BD Difco), and cation-adjusted MHB (BD Difco) were used throughout. Iron-depleted cation adjusted Mueller-Hinton broth (ID-CA-MHB) was used for experiments with cefiderocol and prepared according to CLSI guidelines^37^: CA-MHB was treated with 1% chelex resin (Sigma-Aldrich) for 2 h at room temperature with gentle stirring, filtered to remove chelex, and 11.25 µg/mL MgCl_2_, 22.5 µg/mL CaCl_2_, and 10 µM ZnCl_2_ were added back. Cefiderocol stock solution for experiments was prepared by incubating cefiderocol 30 µg disks (Hardy) with water at 37°C with shaking for 1 h to elute cefiderocol into solution at 420 µg/mL final concentration and subsequently filter sterilized, or by creating 10 mg/mL stock solution in DMSO of cefiderocol powder (MedChemExpress). Either stock preparation produced comparable results. For broth cultures, bacteria were cultured at 37°C with shaking.

### Population analysis profile

Population analysis profile (PAP) was performed as described previously^17^ and indicated in Figure S1. A given clinical isolate was grown overnight from a single colony streaked to MHA from -80°C glycerol stocks in 1.5 mL ID-CA-MHB. After approximately 16-20 h of growth, the culture was serially diluted in PBS in a 96-well plate (Falcon) and 7.5 or 10 µl of each dilution was plated on MHA containing antibiotics as indicated. Colonies were enumerated after 24-48 h of growth. Isolates were classified as resistant if the number of colonies that grew at the breakpoint concentration was at least 50% of those that grew on antibiotic free plates. If the isolate was not resistant, it was classified as heteroresistant if the number of colonies that grew at 2 or 4 times the relevant breakpoint was at least 0.0001% (1 in 10^6^) of those that grew on antibiotic free plates. If the isolate was neither classified as resistant nor heteroresistant, it was classified as susceptible. The limit of detection is approximately -7 logs, in the figures the y-axis is set at -6 logs of survival. See Figure S1 for graphical representation.

In defining the features of heteroresistance in the isolates of this work, based on four guidelines set forth by Andersson et al^18^:

1. Clonality: these isolates demonstrate monoclonal heteroresistance, they are purified isolates and single colonies are used throughout experiments
2. Level of resistance: the MIC of the resistant subpopulation in these strains is ≥8X the MIC of the main population, when comparing the amount of killing by the lowest concentration of cefiderocol used throughout, and the growth of the resistant subpopulation at 32 µg/mL.
3. Frequency of the resistant subpopulation: We consider the frequency of the subpopulation at 2X the CLSI breakpoint, for the strains used throughout, the frequency is ∼0.01% to 0.001%.
4. Stability: these isolates demonstrate unstable heteroresistance. As shown in the passage experiments after selection, there is a significant reduction in the resistant population frequency within 50 generations (∼10 generations per passage) of growth in antibiotic free media.

### Time kill

RS was streaked from -80 glycerol stocks to MHA for isolation and a single colony was used to start overnight cultures in ID-CA-MHB in 1.5 ml volume in 5 ml aeration culture tubes and grown for 10 h. Subsequently, 6 µl of this overnight culture was added to 3 ml ID-CA-MHB containing 16 µg/mL cefiderocol in 10 ml aeration culture tubes and grown for 24 h. At the timepoints indicated, 100 µl of culture was removed, serially diluted in PBS and plated on MHA containing 0 or 2 µg/mL cefiderocol, grown for 24 h and surviving colonies were enumerated.

### Resistance stability assays

Strains were streaked from -80 glycerol stocks to MHA for isolation and a single colony was used to start overnight cultures in ID-CA-MHB in 1.5 ml volume in 5 ml aeration culture tubes and grown for 15-20 h. Cultures were back-diluted into 3 ml of fresh ID-CA-MHB in 10 ml aeration culture tubes and grown with cefiderocol for 24 h for RS and Mu1984 and for 48 h for Mu1956 or without cefiderocol for 24 h for all strains. The conditions for each strain are listed in Materials and Methods Table 1. Aliquots were taken to determine the percent resistance to cefiderocol to ensure selection of the resistant subpopulation (see Materials and Methods Table 1). The remaining cells were collected and gDNA was extracted with Promega Wizard Genomic Extraction kit for whole genome sequencing. To confirm the resistant population selected in the presence of cefiderocol was the result of heteroresistance and not a spontaneous resistance mutation, the culture grown in cefiderocol was passaged each day by 1:1000 back-dilution into fresh ID-CA-MHB and grown for 24 h, with aliquots taken to quantify the resistant population and determine if the frequency substantially reduced after removal of antibiotic.

### Whole genome sequencing

The samples from the resistance stability assays were subject to whole genome sequencing. The genome of the strains grown without antibiotic (mostly susceptible) was subjected to Illumina (650 Mbp) and Nanopore (300 Mbp ONT) sequencing to create a reference genome and quantify gene copy variation. The genome of the resistant population from growth with cefiderocol was subject to Illumina sequencing and mapped to the respective reference genome with CNV analysis. Quality control and adapter trimming was performed with bcl2fastq^38^. Reads were mapped to their respective references via bwa mem^39^. PCR and optical duplicates were marked and excluded from the analysis using PicardTools’ ‘MarkDuplicates’^40^ functionality. Aligned read counts were imported into R’s CNOGpro^41^package. CNV events were called via CNOGpro using a bootstrapping method, which calculates an average gene number event giving a possible upper and lower bound. The new reference genomes have been uploaded to NCBI (see Materials and Methods Table 1). Genome sequencing and analysis were performed by Microbial Genome Sequencing Center (migscenter.com) and SeqCenter (seqcenter.com).

### *Enterobacter* RS mutagenesis

Lambda-red based allelic exchange was used to replace the amplicon or the *blaSHV-5* coding sequence with a kanamycin resistance gene, and then the gene was removed with Flp recombinase^42,43^ to create unmarked in-frame deletions. The kanamycin resistance gene from pEXR6K_*kanFRT* was cloned using Promega GoTaq 2X master mix with Flp recognition sequence and homology to regions flanking the gene to be replaced using primers JCP293/244 for *blaSHV-5* and JCP269/271 for the amplicon region (Materials and Methods Table 2). The purified PCR product was electroporated into competent RS pKD46-tet and transformants were selected on 90 µg/mL kanamycin. Transformants were re-streaked to Kan90 for isolation and subject to PCR for successful allelic exchange as indicated by a product size change using primers flanking the gene to be replaced: JCP213/214 for *blaSHV-5* and JCP272/210 for the amplicon region (Materials and Methods Table 2). The mutants were then made electrocompetent and electroporated with pCP20^43^. Transformants were selected at 30°C on 50 µg/mL chloramphenicol, patched to 50 µg/mL chloramphenicol at 30°C and grown for 24 h, then patched to MHA and MHA+Kan90. Kanamycin-sensitive mutants were streaked for isolation and PCR was used with the same flanking primers to screen for the loss of the kanamycin resistance gene.

### *Enterobacter* RS *blaSHV-5* manipulation and allelic exchange

To manipulate *blaSHV-5,* the region of the chromosome containing the coding sequence, and 750 bp upstream and 747 bp downstream were cloned using primers JCP343/344 into SmaI (NEB)-digested pTOX5 allelic exchange vector^44^, using *E. coli* PIR2 and sequenced to confirm using primers JCP345/346/347/348 (GeneWiz). PIR2 carrying pTOX5 plasmids was propagated in MHB or MHA containing 20 μg/mL chloramphenicol and 2% glucose. The wildtype gene, the C-terminal myc-epitope tagged, the S238G;K240E variant, and the M69I variant was used to complement the Δ*blaSHV-5* deletion. The myc epitope was added before the stop codon of *blaSHV-5* by NEB Q5 Site-Directed Mutagenesis according to the manufacturer using primers JCP505/506 with pTOX5 *blaSHV-5* template. S238G;K240E and M69I refers to the Ambler numbering system for beta-lactamases^45^, in RS SHV-5 the numbering is S236G;K237E and M67I. The S238G;K240E mutations were created using the NEB Q5 Site-Directed Mutagenesis reagents according to the manufacturer using primers JCP323/324 and subsequently JCP325/326. M69I was created using primers JCP321/322. The resulting plasmids were sequenced with primers JCP213, JCP214, JCP345, and JCP346. The pTOX5 *blaSHV-5* vectors were transformed into electrocompetent *Enterobacter* RS Δ*blaSHV-5* with 25 µg/mL chloramphenicol and 2% glucose. Purple colonies were selected, resuspended in 2 ml LB+2% glucose, cells were collected when OD of culture reached approximately OD 0.2, the cells were washed twice in M9 salts + 2% rhamnose, and resuspended in 500 μl of M9 salts + 2% rhamnose. 200 μl and 20 μl were each plated on M9 agar+2% rhamnose and grown at 37°. After 36-48 h, non-purple colonies were restreaked to MHA. Allelic exchange was confirmed by PCR using primers JCP213/214, and primers JCP255, JCP256, and JCP258 were used for Sanger sequencing of JCP213/214 PCR product (GeneWiz). The chromosomal loci of each strain were 100% nucleotide match to the original chromosome except for introduced mutations.

### RS confirmation of duplication

PCR was performed using GoTaq Master Mix (Promega) using 5 ng of DNA extracted from RS collected after growth on MHA containing 0 or 32 μg/mL cefiderocol, with 30 cycles of amplification. Primer pair 1 shown in Figure S4 is JCP255/238 and primer pair 2 is JCP237/229 (Materials and Methods Table 2). An equal volume of PCR product was loaded into 0.7% agarose containing ethidium bromide and imaged using Biorad Chemidoc XRS+ with Image Lab 6.1 software.

### Quantitative PCR of DNA

qPCR was performed from samples as described in each figure legend. DNA was purified using the Wizard Genomic DNA Purification kit (Promega) following the manufacturer’s instructions. DNA was diluted to 10 ng/μl following quantification by Take3 (Biotek). Power SYBR Green PCR Master Mix (Applied Biosystems, Thermo Fisher) was used in 20 μl reactions with 10 ng of DNA and 50 nM primers. For each gene, multiple primer pairs were tested in advance of use according to guidelines of Applied Biosystems StepOnePlus Real-Time PCR machine to confirm primer pairs had similar efficiencies and melt curves. qPCR was performed in technical triplicate and fold change was calculated from the mean of technical triplicates. C_T_ values were normalized to the respective housekeeping gene. For all experiments except where noted, the fold change for each biological replicate was compared to the same replicate from no antibiotic conditions, resulting in a value of 1 for each no antibiotic replicate. Fold change was calculated using the 2^-ΔΔCT^ method^46^. Primers are listed in Materials and Methods Table 2.

RS was grown overnight in 1.5 ml ID-CA-MHB, serially diluted in sterile PBS, and 7.5-10 μl of each dilution were spot-plated on MHA. After ∼24 h of growth at 37°, cells were collected. Surviving colonies from MHA containing no addition or various cefiderocol and potassium clavulanate (Combi-blocks) concentrations (as described in Figures) was collected with sterile cotton swab and resuspended in PBS, then cells were collected by centrifugation. For qPCR, *cysG* was detected with JCP241/242 and was used as the housekeeping gene for normalization, *blaSHV-5* with JCP257/258, NF29_02980 with JCP253/254, *recF/recN/SMC* (NF29_03000) with JCP251/252, *aldolase* (NF29_03030) with JCP237/238, and NF29_03450 with JCP265/266.

Mu1956 was grown overnight in 1.5 ml ID-CA-MHB and 12 μl were subcultured into 3 ml ID-CA-MHB alone or with addition of 8 μg/mL cefiderocol, or addition of 8 μg/mL cefiderocol and 0.5 μg/mL avibactam sodium (Combi-blocks), in a 10 ml volume aeration tube grown at a 45° angle with shaking. After 20 h, an aliquot was serially diluted and plated to MHA with and without cefiderocol addition and the rest of the cells were collected by centrifugation for DNA extraction. For qPCR *clpX* was detected with JCP302/303 and was used as the housekeeping gene for normalization, *blaADC-30* was detected with JCP298/299.

Mu1984 was grown overnight in 1.5 ml ID-CA-MHB and 12 μl were subcultured into 3 ml ID-CA-MHB with no addition or containing 8 μg/mL cefiderocol, 8 μg/mL cefiderocol and 0.5 μg/mL avibactam, in a 10 ml volume aeration tube grown at a 45° angle with shaking. After 20 h, an aliquot was serially diluted and plated to MHA with and without cefiderocol addition and the rest of the cells were collected by centrifugation for DNA extraction. For this strain, selection was less consistent and the samples for which <50% of the population grew on MHA containing 2 μg/mL cefiderocol were removed from analysis. For qPCR *clpX* was detected with JCP302/303 and was used as the housekeeping gene for normalization*, blaADC-33* was detected with JCP337/338.

The metallo-beta-lactamase producing *K. pneumoniae* was grown overnight in 1.5 ml ID-CA-MHB, serially diluted in sterile PBS, and 7.5-10 μl of each dilution were spot-plated on MHA containing 0 or 16 μg/mL cefiderocol. After ∼24 h of growth at 37°, surviving colonies was collected with sterile cotton swab and resuspended in PBS, then cells were collected by centrifugation. For qPCR, *rplS* was detected with TOP54/55 and was used as the housekeeping gene for normalization, *blaNDM-5* with TOP52/53.

### qPCR for mRNA, qPCR for DNA, and immunoblot of RS Δ*blaSHV-5*::*blaSHV-5* and RS Δ*blaSHV-5*::*blaSHV-5_myc*

RS strains were grown overnight in 1.5 ml ID-CA-MHB, diluted in sterile PBS 1 in 10^-6^ for MHA and 1:5 for MHA+ 32 µg/ml cefiderocol, and 200 µl were spread. After ∼24 h of growth at 37°, cells were collected in sterile PBS and 1:1:4 (volume) aliquots of the suspension were collected by centrifugation. qPCR for *blaSHV-5* gene abundance was performed as above.

For qPCR to quantify blaSHV-5 mRNA transcript abundance, the cell pellet was treated with RNAProtect (Qiagen) and RNA was isolated with Qiagen RNAEasy (#74104), then treated with DNAse (BioLab #M0303S) according to manufacturer’s instructions. Reverse-transcriptase quantitative PCR was performed with Power SYBR Green RNA to C_T_ 1-Step Kit (Applied Biosystems #4389986). *cysG* was detected with JCP241/242 and was used as the housekeeping for normalization, *blaSHV-5* with JCP257/258. For all experiments except where noted, the fold change for each biological replicate was compared to the same replicate from no antibiotic conditions, resulting in a value of 1 for each no antibiotic replicate. Fold change was calculated using the 2^-ΔΔCT^ method^46^.

For immunoblot, the cell pellet was resuspended with B-PER Complete, Bacterial Protein Extraction Reagent (#89821) with a Pierce Protease Inhibitor tablet, EDTA free (#A32965), incubated for 15 minutes, and centrifuged. The soluble fraction was collected and protein content quantified with BCA assay (Thermo). Twenty µg of protein in Laemmli buffer was loaded for each sample and subjected to SDS-PAGE, transferred to nitrocellulose membrane and stained with Ponceau. After washing the membrane was probed with 1:500 anti-myc (Cell Signaling #2276; 9B11) followed by 1:15,000 IRDye 680RD (donkey anti-mouse; Li-Cor#926-68072) and imaged with Li-Cor Odyssey FC.

### SHV-5 enzyme kinetics

Primers JCP501/502 were used to clone blaSHV-5 from the RS chromosome and assembled with pET28a-TEV (Palzkill laboratory) amplified with JCP523/524 using Hifi assembly, resulting in hexa-His, TEV site, and SHV-5 lacking the signal sequence. Sequencing with commercial T7 and T7-Term primers confirmed the constructs (Genewiz). The plasmid was transformed into NEB5α with 35 μg/mL kanamycin selection and transformed into *E. coli* BL21(DE3) cells for expression.

#### Enzyme Expression

*E. coli* BL21(DE3) cells with the pET28a-*blaSHV-5* plasmid were cultured in lysogeny broth medium at 37°C until the OD_600_ reached 0.8 to 1.0. Protein expression was initiated using 0.2 mM isopropyl β-D-1-thiogalactopyranoside (IPTG). The culture was then placed in a shaking incubator set at 23°C for 20 h. The next day, the cells were pelleted using low-speed centrifugation, washed with phosphate-buffered saline (PBS), and stored at -80°C.

#### Enzyme Purification

The cells were thawed and resuspended in binding buffer (25 mM sodium phosphate, pH 7.4, 300 mM NaCl, 20 mM imidazole, and Xpert protease inhibitor cocktail solution [GenDEPOT]). The cells were then lysed using sonication, and the cell debris was pelleted by centrifugation at 10,000 x g for 15 min at 4°C. The supernatant was filtered through a 0.45 µm filter. The Cobalt Chelating Resin (G-Biosciences) was washed with water and preequilibrated with binding buffer. The supernatant and preequilibrated resin were mixed for 1 hr at 4°C. The mixture was then loaded onto a gravity flow column. The flowthrough was collected and poured over the resin for a total of three times. The resin was washed with binding buffer, and the protein was eluted by adding buffer (25 mM sodium phosphate, pH 7.4, 300 mM NaCl, and Xpert protease inhibitor cocktail solution) supplemented with 40, 60, 80, 100, 150, and 200 mM imidazole, respectively. The presence and purity of SHV-5 in each fraction were visualized by sodium dodecyl sulfate-polyacrylamide gel electrophoresis (SDS-PAGE) followed by Bio-Safe Coomassie G-250 staining. Protein fractions containing SHV-5 were pooled, concentrated, and buffer-exchanged with wash buffer (25 mM sodium phosphate, pH 7.4, 300 mM NaCl, and Xpert protease inhibitor cocktail solution) using an Amicon Ultra-15 centrifugal filter unit (MilliporeSigma). To remove the His-tag, the protein was incubated with tobacco etch virus (TEV) protease overnight at 4°C. The next day, Ni Sepharose 6 Fast Flow Resin (Cytiva) was applied to the reaction mixture to remove the His-tagged TEV protease and any remaining SHV-5-His. Protein purity and His-tag cleavage were visualized by SDS-PAGE followed by Coomassie staining. We found that the SHV-5 M69I enzyme was purified at low yield, possibly due to reduced stability relative to SHV-5. Due to the lower yield, we were not able to purify the TEV-protease cleaved version with the His-tag removed in sufficient quantity. Therefore, the His-tag version of SHV-5 M69I was used for enzyme kinetics assays. We also showed that the His-tagged version of SHV-5 has similar kinetic parameters as SHV-5 without the His-tag, showing that the His-tag does not affect enzyme function.

#### Kinetic assays

Michaelis-Menten steady-state kinetic parameters for SHV-5 were determined for cefiderocol, cephalothin, and cefotaxime. The wavelengths and extinction coefficients used were the following: cefiderocol, 259 nm, Δε = 9443 M^-1^ cm^-1^; cephalothin, 262 nm, Δε = 7660 M^-1^ cm^-1^; cefotaxime, 264 nm, Δε = 7250 M^-1^ cm^-1^. Reactions for each substrate were conducted at 25°C in buffer containing 50 mM sodium phosphate (pH 7.0) and 1 µg/mL bovine serum albumin. A Beckman Coulter DU 800 spectrophotometer (Beckman Coulter) was used to monitored substrate hydrolysis. Initial hydrolysis rates were plotted against the substrate concentrations, and the data was fitted to the Michaelis-Menten equation using GraphPad Prism 10 (GraphPad Software, www.graphpad.com) to determine *k*_cat_ and *K*_M_ values. The error for *k*_cat_/*K*_M_ values was calculated using the equation below where SEM is the standard error of the mean:

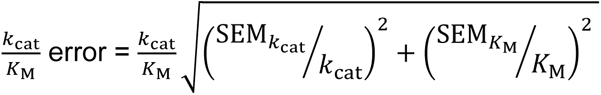

### Galleria mellonella infection model

*G. mellonella* larvae with an average mass of ∼200 mg were used within two weeks of delivery from Speedyworm.net. Larvae were maintained in ambient air at ∼15°C until time of experiment. The agar used was MacConkey Agar with Crystal violet, sodium chloride and 0.15% bile salts (MACVBS) containing 1 μg/mL ciprofloxacin (total CFU) or 1 μg/mL ciprofloxacin + 2 μg/mL cefiderocol. *Enterobacter* RS is ciprofloxacin resistant, and MACVBS +ciprofloxacin prevents the growth of any larvae associated bacteria other than RS. Like MHA containing 2 μg/mL cefiderocol, MACVBS containing no antibiotic, ciprofloxacin, or cipfloxacin + 2 μg/mL cefiderocol resulted in no change in *blaSHV-5* abundance. *Enterobacter* strain RS was grown overnight in Mueller-Hinton Broth at 37°C with shaking. Cells were collected by centrifugation, washed twice in sterile PBS, and diluted in PBS to a density of ∼1×10^8^ CFU/mL. 10 μl of the bacterial suspension was inject with a sterilized Hamilton #1701 Gastight 10 µl syringe and worms were moved to 37°C. The inoculum was serially diluted in PBS and plated on MACVBS+ ciprofloxacin; after 24 h, the inoculum density was enumerated and CFU collected for qPCR. At 1 and 4 hours post infection, worms were injected with 10 μl PBS or 10 μl cefiderocol solution in PBS (0.5 mg/mL) to achieve 25 mg/kg at each dose. At ∼20 hours post infection, each worm was added to 1 mL sterile PBS and homogenized using Biospec Products Tissue Tearor #985370-395. The homogenate was serially diluted in PBS and 5 μl of each dilution was spot plated on agar, and 200 μl of undiluted homogenate was spread on agar as well. After 24 h, colony forming units (CFU) were enumerated to calculate the total CFU and CFU on agar containing 2 μg/mL cefiderocol. The resistant CFU were collected for qPCR. qPCR for *blaSHV-5* abundance was performed as described above, with the inoculum as the comparator control for calculating ΔΔC_T_.

### Beta-lactamase Inhibitor screen

To screen carbapenem-resistant isolates with beta-lactamase inhibitors, the cefiderocol PAP was performed as above, except the MHA contained cefiderocol alone or with either 4 µg/mL clavulanate or 4 µg/mL avibactam. Each inhibitor was also included in MHA with no cefiderocol to confirm that the inhibitor alone did not affect total colony forming units.

### ddPCR

Droplet digital PCR was completed at the Emory Integrated Genomics Core using the Bio-Rad QX200 ddPCR system using the QuantaSoft Analysis Pro Software. *blaSHV-5* primers were JCP257/258, and the custom probe internal to the PCR product was conjugated to FAM by Bio-Rad (#10031276) and cysG primers were JCP241/242, the custom prober internal to the PCR product was conjugated to HEX (#10031279), Black Iowa was the quencher. The custom internal probe for blaSHV-5 was GATCTGGTGGACTACTCGCC, and for cysG: GGGCAATCTGACCAAGGTGC. The parameters were: 95°C for 10 min, 40 cycles of (94°C for 30 s, 60°C for 1 min), 98°C for 10 min, hold at 4°C. 100 ng of DNA was used in each well.

### RS SHV-5 variant experiments

PAP was performed as described above but with cefiderocol from 0.125 µg/mL to 32 µg/mL for RS mutant strains: Δ*blaSHV-5,* Δ*blaSHV-5*::*blaSHV-5* (referred to as *blaSHV-5* in Figure 3), Δ*blaSHV-5*::*blaSHV-5* M69I (*blaSHV-5* M69I), and Δ*blaSHV-5*::*blaSHV-5* S238G;K240E (*blaSHV-5* S238G;K240E; which is SHV-1). Surviving colonies were collected from MHA containing 0 or 4 µg/mL cefiderocol and suspended in 1 mL sterile PBS, used for DNA extraction, and qPCR was performed as described above. 10 µl of the suspension was added to 1 mL ID-CA-MHB containing 4 µg/mL cefiderocol. After 24 h growth in broth containing 4 µg/mL cefiderocol, PAP was performed. After 24 h of growth, surviving colonies were again collected from MHA containing 0 (this represents the gene abundance after growth in 4 µg/mL cefiderocol in Figure 3 panels C, F, and I) or 8 µg/mL cefiderocol. From the suspension of cells collected from 8 µg/mL cefiderocol, 10 µl of the suspension was added to 1 mL ID-CA-MHB containing 8 µg/mL cefiderocol. After 24 h growth in broth containing 8 µg/mL cefiderocol, PAP was performed. After 24 h of growth, surviving colonies were again collected from MHA containing 0 (this represents the amplification number after growth in 8 µg/mL cefiderocol in Figure 3 panels C, F, and I) or 16 µg/mL cefiderocol. From the suspension of cells collected from 16 µg/mL cefiderocol, 10 µl of the suspension was added to 1 mL ID-CA-MHB containing 16 µg/mL cefiderocol. After 24 h growth in broth containing 16 µg/mL cefiderocol, PAP was performed. After 24 h of growth, surviving colonies were again collected from MHA containing 0 (this represents the amplification after growth in 16 µg/mL cefiderocol in Figure 3 panels C, F, and I) or 32 µg/mL cefiderocol and was used for DNA extraction and qPCR was performed (Figure 3 panels E, H, and K). For qPCR, the surviving colonies collected from MHA containing no cefiderocol in the initial PAP is the comparator for calculating ΔΔC_T._

### RS cefiderocol/clavulanate passage

PAP was performed as described above on cefiderocol with 4 µg/mL clavulanate, as well as plated on MHA containing 8 µg/mL cefiderocol and 0.06 µg/mL clavulanate. After ∼24 h of growth, surviving colonies were enumerated for each replicate and collected from MHA containing no cefiderocol (the baseline comparator for qPCR). The surviving colonies from MHA containing 8 µg/mL cefiderocol and 0.06 µg/mL clavulanate were resuspended in 1mL PBS from which 7.5 µl was inoculated into 2 mL ID-CA-MHB with 8 µg/mL cefiderocol and 0.06 µg/mL clavulanate and the remaining cells were collected for DNA extraction. After 24 h of growth, PAP was performed on cefiderocol with 4 µg/mL clavulanate and the culture was also plated on MHA containing 16 µg/mL cefiderocol and 0.25 µg/mL clavulanate. After 24 h, surviving colonies were enumerated and collected from MHA containing 16 µg/mL cefiderocol and 0.25 µg/mL clavulanate and resuspended in 1ml PBS. 7.5 µl was added to 2 mL ID-CA-MHB with 16 µg/mL cefiderocol and 0.25 µg/mL clavulanate and the remaining cells were collected for DNA extraction. After 24 h of growth, PAP was performed on cefiderocol with 4 µg/mL clavulanate and the culture was also plated on MHA containing 8 µg/mL cefiderocol and 4 µg/mL clavulanate. After 24 h, surviving colonies were enumerated and collected from 16 µg/mL cefiderocol and 4 µg clavulanate for qPCR.

### Mu1956 passage experiment

Mu1956 was grown overnight and PAP performed on MHA containing 0-32 µg/mL cefiderocol and 4 µg/mL avibactam. From the overnight culture, 12 µl were inoculated into 3 ml ID-CA-MHB without drug or with 4 µg/mL cefiderocol and 1 µg/mL avibactam, in a 10 ml volume aeration tube grown at a 45° angle with shaking. The cells of the overnight broth culture collected for qPCR. After 24 h, PAP was performed on MHA containing cefiderocol and 4 µg/m avibactam. The cells of the broth cultures were collected for qPCR.

### Broth Microdilution

Broth microdilution was used to determine the minimum inhibitory concentrations (MICs) of cefiderocol for strains grown in ID-CAMHB with and without cefiderocol by following the CLSI protocol^23^. RS was grown for 24 h in 0 or 32 µg/mL cefiderocol, and Mu1984 and Mu1956 were grown for 24 h in 0 or 8 µg/mL cefiderocol. In brief, a suspension of bacteria was diluted in ID-CA-MHB to 5×10^4^ CFU/well; cefiderocol concentrations were prepared in 2-fold dilutions ranging from 0.0125 to 64 µg/mL. After incubation of 16 hours (or 20 hours for *A. baumannii*) at 37°C, MICs were determined according to the well with the lowest concentration in which bacterial growth was not visible.

### Statistical Methods

Figure legends detail the n of each experiment, statistical test employed where relevant, and definition of the error bars.

### Graphics

Some graphical content was from BioRender.com (2022), used with Publication Rights under agreements TK256AOOMU and RH256AOS93.

**MATERIALS AND METHODS TABLE 1.**
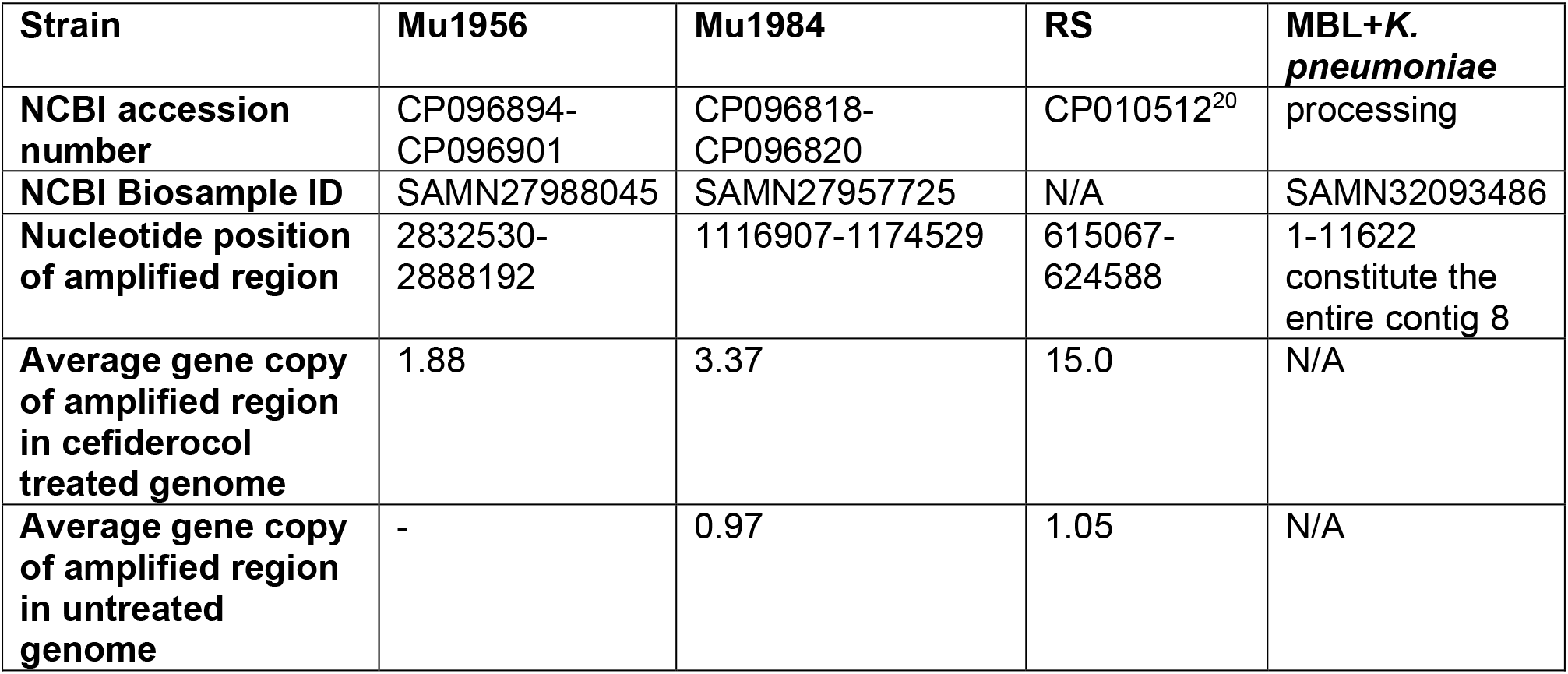
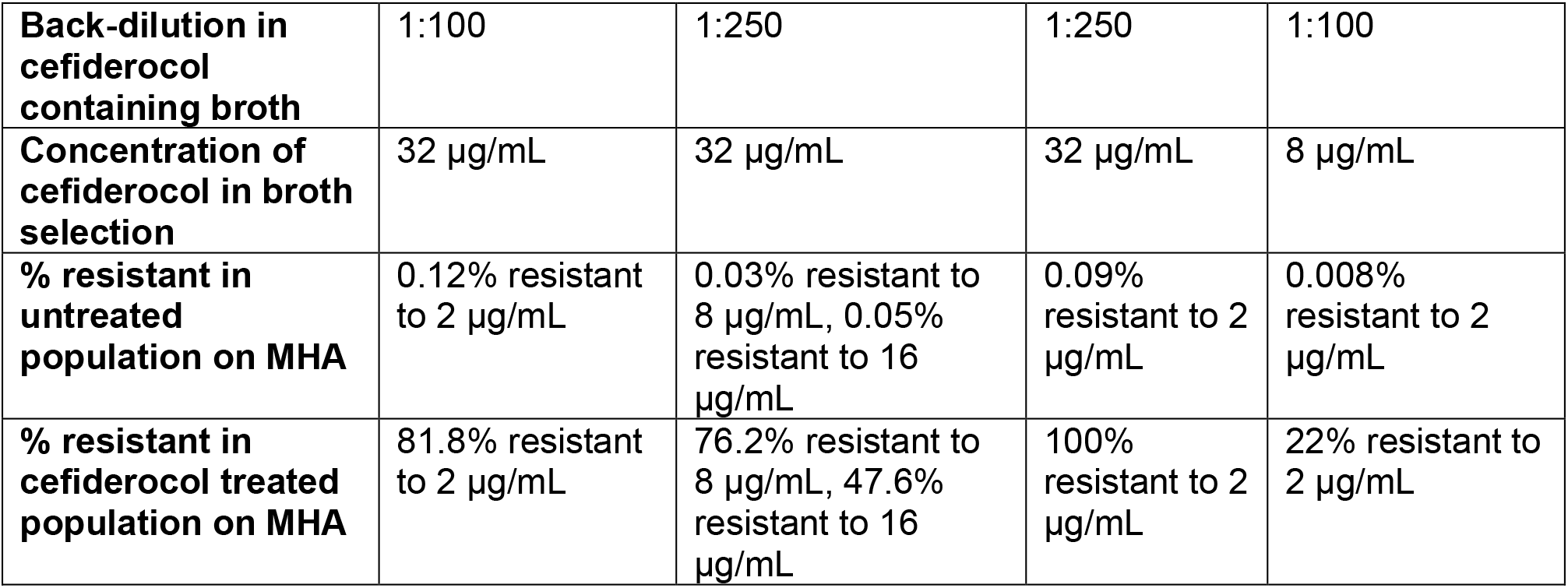
Genome sequencing information.

**MATERIALS AND METHODS TABLE 2.**
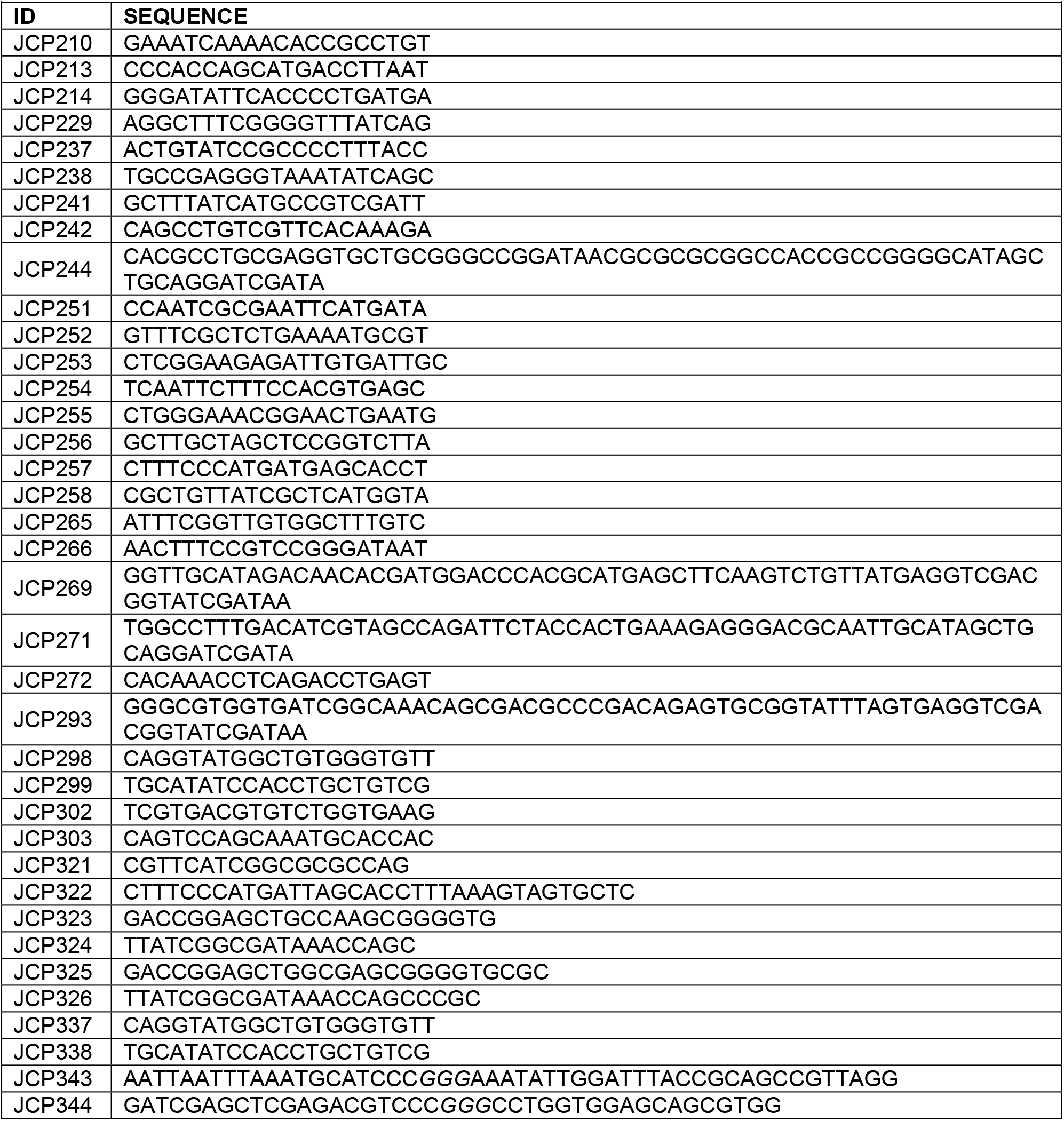
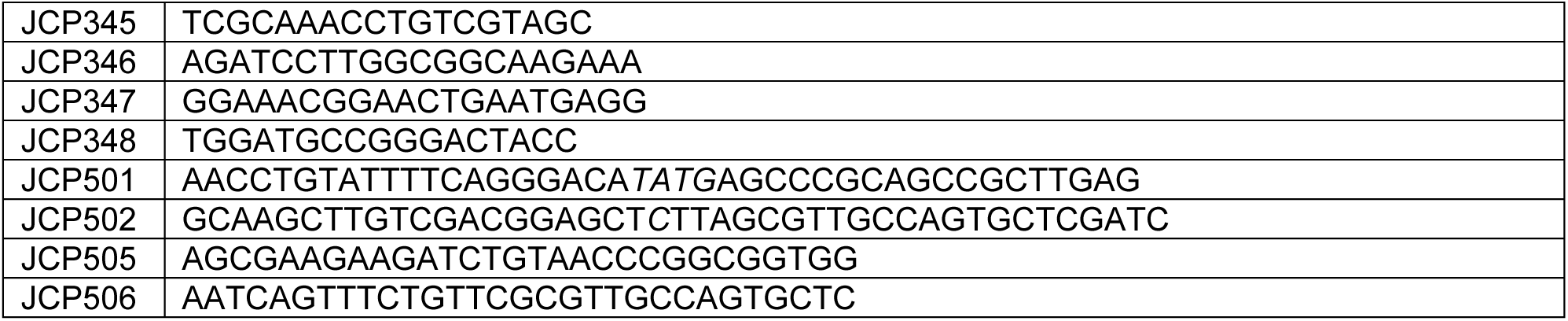
PRIMERS.

## AUTHOR CONTRIBUTIONS

JEC: Conceptualization, Investigation, Project Administration, Visualization, Writing-original draft.

TO: Investigation, Validation, Writing-Review & Editing

CNA: Investigation, Validation, Writing-Review & Editing

CN: Investigation, Writing-Review & Editing

JMC: Investigation, Writing-Review & Editing

SWS: Resources, Writing-Review & Editing

PNR: Supervision, Resources, Writing-Review & Editing

TP: Supervision, Writing-Review & Editing

DSW: Conceptualization, Project Administration, Supervision, Funding Acquisition, Writing-Review & Editing

## MATERIALS AND CORRESPONDENCE

Genome sequences are available at NCBI Genbank with accession numbers CP096894-CP096901 (Mu1956), CP096818-CP096820 (Mu1984), and NZ_JAQGEE010000000 (MBL-producing *K. pneumoniae*). Material requests should be made to David S. Weiss.

## ACKNOWLEDGEMENTS

We would like to thank the US Centers for Disease Control and Prevention (CDC)’s Emerging Infections Program and the Georgia Multi-Site Gram-negative Surveillance Initiative (MuGSI), including Jesse Jacob and Monica Farley, for providing isolates. We acknowledge the contribution of genome sequencing by Microbial Genome Sequencing Center (MiGS) and SeqCenter. This study was supported in part (ddPCR) by the Emory Integrated Genomics Core (EIGC) (RRID:SCR_023529), which is subsidized by the Emory University School of Medicine and is one of the Emory Integrated Core Facilities. Additional support was provided by the Georgia Clinical & Translational Science Alliance of the National Institutes of Health under Award Number UL1TR002378.This work was supported by the National Institutes of Health [AI158080, AI141883, AI32956] and the Department of Veteran’s Affairs [BX002788], and the National Institutes of Health’s Office of the Director, Office of Research Infrastructure Programs, P51 OD011132. DSW is supported by a Burroughs Wellcome Fund Investigators in the Pathogenesis of Infectious Disease award and JEC is supported by the Cystic Fibrosis Foundation and National Institutes of Health [T32 DK108735]. The Multi-Site Gram-negative Surveillance Initiative is funded by the CDC. The content is solely the responsibility of the authors and does not necessarily reflect the official views of the National Institutes of Health

## SUPPLEMENTAL MATERIAL

**Supplemental Figure 1.**
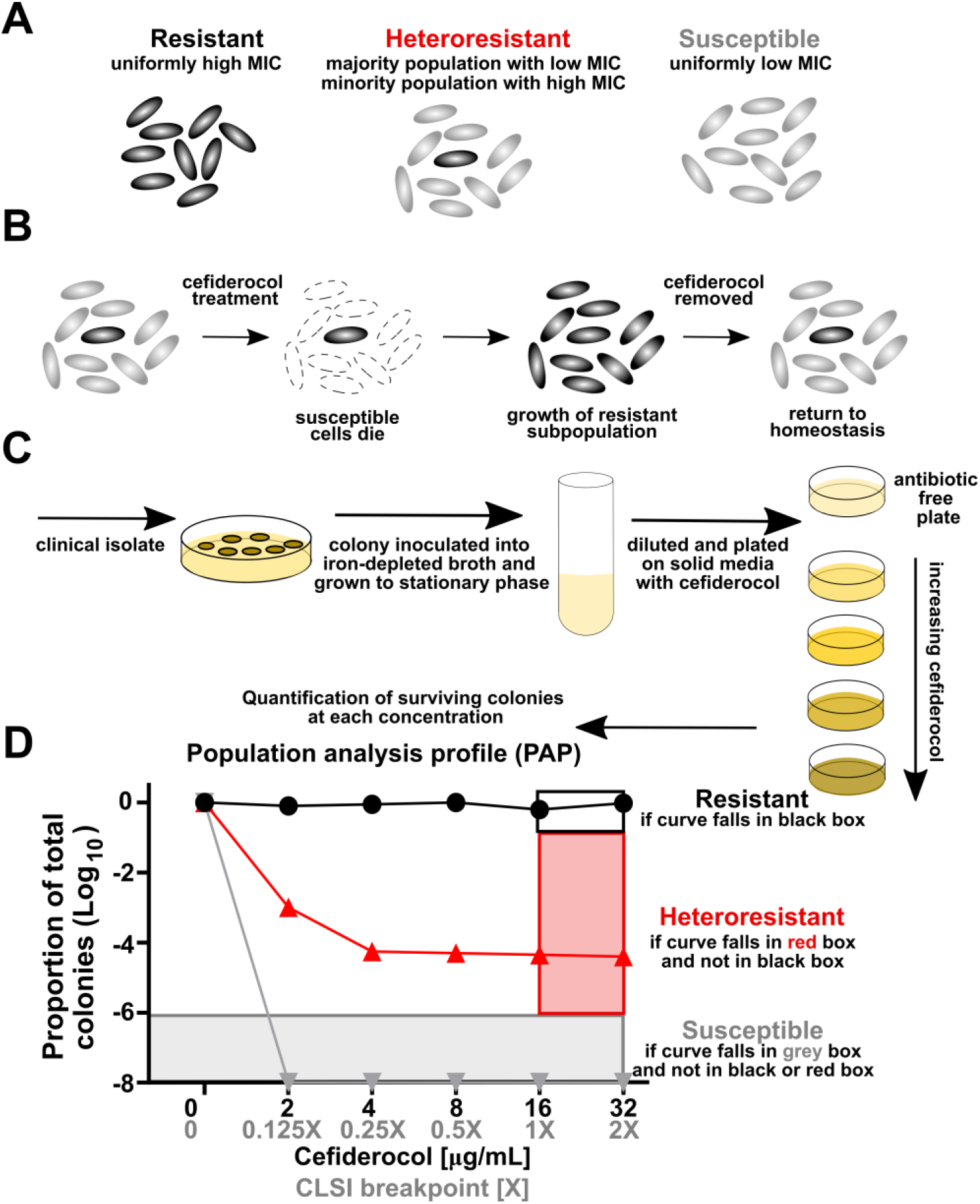
Overview of heteroresistance and population analysis profile (PAP). (**A**) A depiction of the cells grown from a single colony of an isolate exhibiting conventional resistance, heteroresistance, or susceptibility to a given antibiotic. (**B**) Population dynamics of a heteroresistant isolate following cefiderocol treatment. (**C**) Cefiderocol PAP: a clinical isolate is isolated and grown in iron-depleted broth, and then serially diluted, plated, and grown on increasing concentrations of cefiderocol. (**D**) The surviving colonies are enumerated and the isolate is classified as resistant if at least 50% of the total colonies grow at 1 or 2X breakpoint. An isolate is considered susceptible if less than 0.0001% (-6 logs) of the cells grow at any concentration. An isolate is considered heteroresistant if there is less than 50% survival at 1X breakpoint and greater than 0.0001% at 1X and 2X breakpoint. Along the x-axis, the breakpoints based on CLSI are shown in gray, and the corresponding concentration of cefiderocol is in black.

**Supplemental Figure 2.**
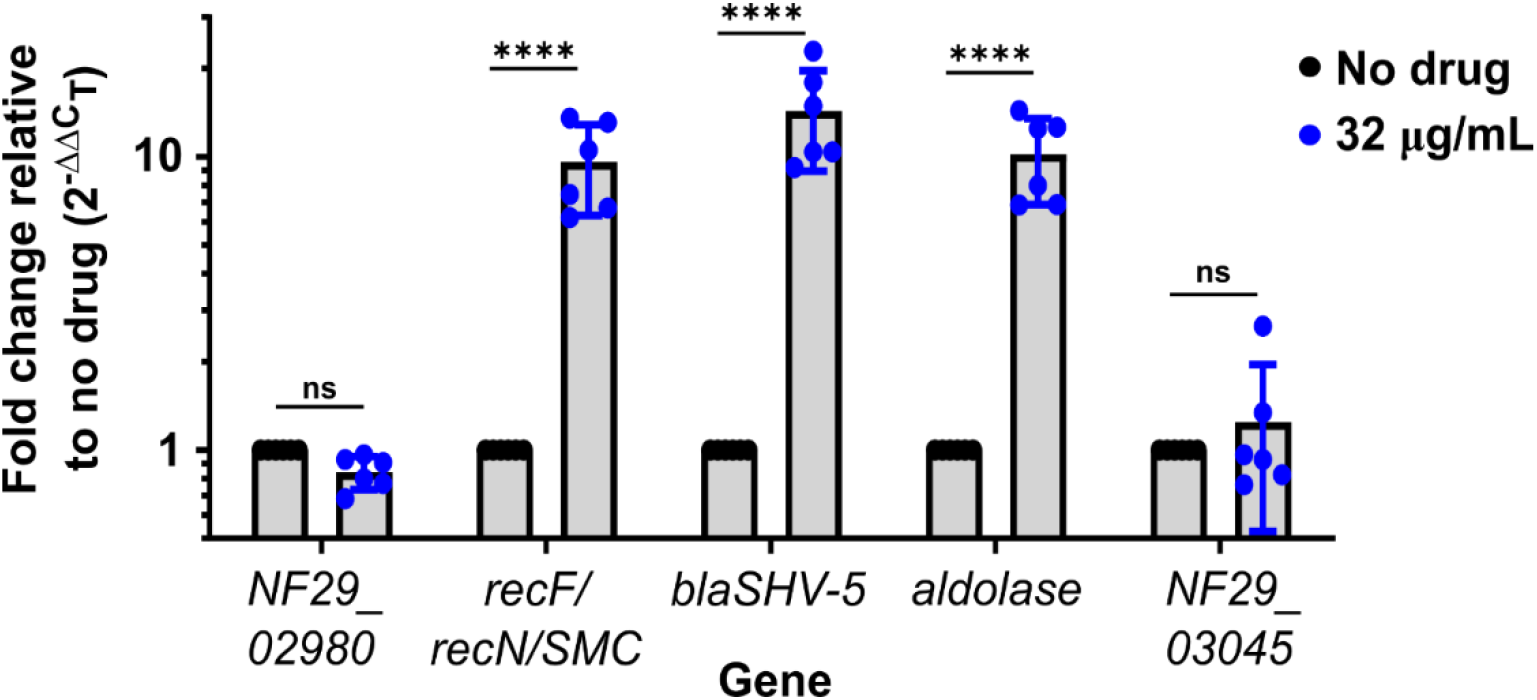
Confirmation of genome sequencing gene abundance variation results by qPCR. Relative abundance of genes outside (*NF29_02980* and *NF29_03045*) and within (*recF/recN/SMC*, *blaSHV-5*, and *aldolase*) the amplified region normalized to *gyrA* (housekeeping gene) as quantified by qPCR from cells collected from Mueller Hinton agar containing no drug or 32 µg/ml cefiderocol. Shown are the means and standard deviation of two independent experiments with 3 biological replicates where each point indicates a biological replicate. **** indicates p<0.0001 by two-way ANOVA, [row factor F (4, 50) = 20.67, column factor F (1, 50) = 114.0] with Sidak’s correction for multiple comparisons.

**Supplemental Figure 3.**
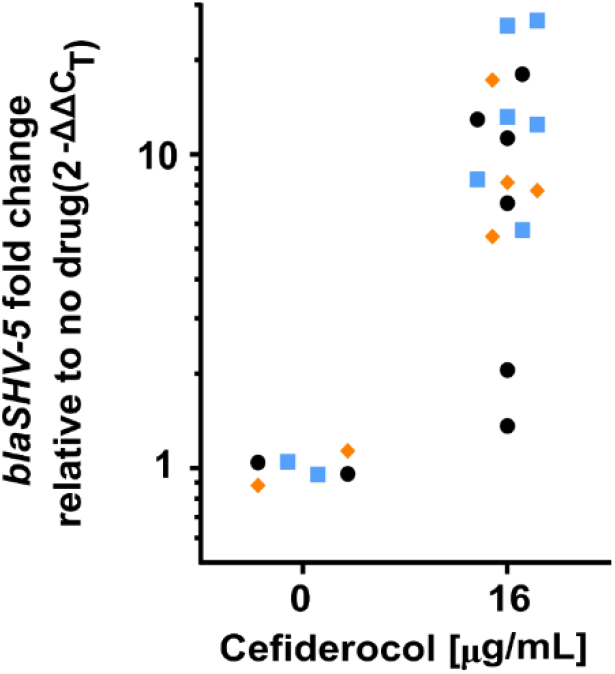
Variation in *Enterobacter* RS *blaSHV-5* gene abundance. Quantification of strain RS *blaSHV-5* abundance normalized to *cysG* by qPCR. An overnight culture was diluted and plated on Mueller Hinton agar alone or containing 16 µg/mL cefiderocol. After growth, a single colony was selected, grown for 7 h in broth without cefiderocol, cells collected, and DNA extracted. The ΔC_T_ of each replicate grown with no drug was averaged to calculate ΔΔC_T._ Each color is a separate biological replicate from one of three unique overnight cultures, and each point is from a unique colony of the respective biological replicate, n=6 for no drug n=16 for 16 µg/mL cefiderocol.

**Supplemental Figure 4.**
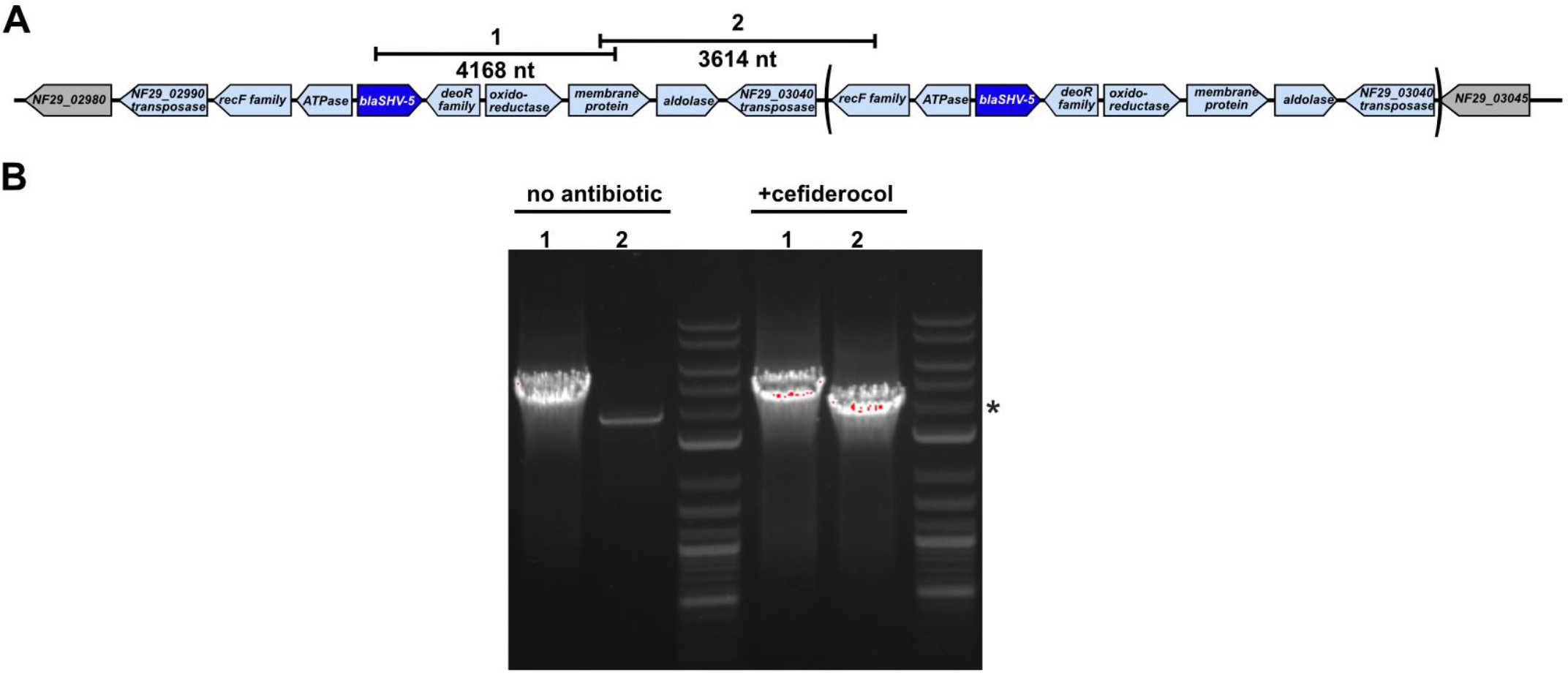
PCR confirmation of *blaSHV-5* amplicon duplication. (**A**) A schematic of the predicted duplication of the *blaSHV-5* region in strain RS. Parentheses indicate duplication of the region with a single intervening copy of the transposase gene. PCR primer pair 1 is internal to the duplicated region, while primer pair 2 spans the duplication event. (**B**) PCR products in agarose gel from primer pairs 1 and 2 using DNA extracted following growth in media alone (first and second lane) or after growth in 32 µg/mL cefiderocol (fourth and fifth lanes) after 30 cycles of PCR. The asterisk indicates the 4 kb marker of the ladders. Note the less abundant product in lane 2 for primer pair 1 after growth in media alone which is consistent with a very small subpopulation of cells exhibiting gene amplification in the absence of cefiderocol.

**Supplemental Figure 5.**
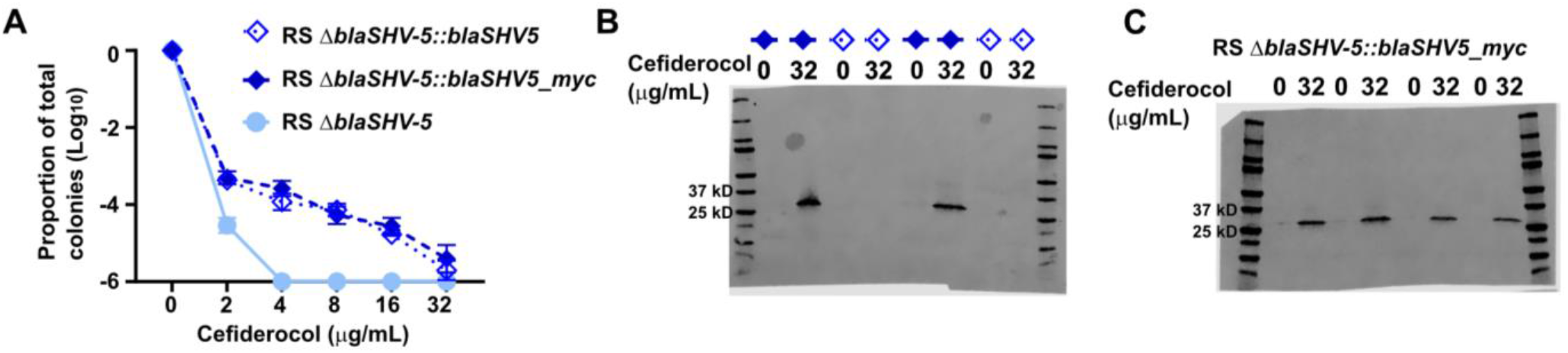
Myc-epitope tagged SHV-5. (**A**) PAP of *Enterobacter* strain RS variants on agar containing cefiderocol. Mean and standard deviation from two independent experiments are shown, from 4-5 biological replicates. (**B**) Anti-myc immunoblot from total cellular lysate from colonies collected from MHA containing 0 or 32 μg/mL cefiderocol of strains RS Δ*blaSHV-5*::*blaSHV-5* and RS Δ*blaSHV-5*::*blaSHV-5_myc*. The reactive band is absent in the strain encoding the native SHV-5. SHV-5_myc is 30.09 kD following cleavage of signal sequence (**C**) Anti-myc immunoblot from RS Δ*blaSHV-5*::*blaSHV-5_myc*. The first three lanes are in Figure 1H. All six replicates of RS Δ*blaSHV-5*::*blaSHV-5_myc* in B-C are quantified in Figure 1I.

**Supplemental Figure 6.**
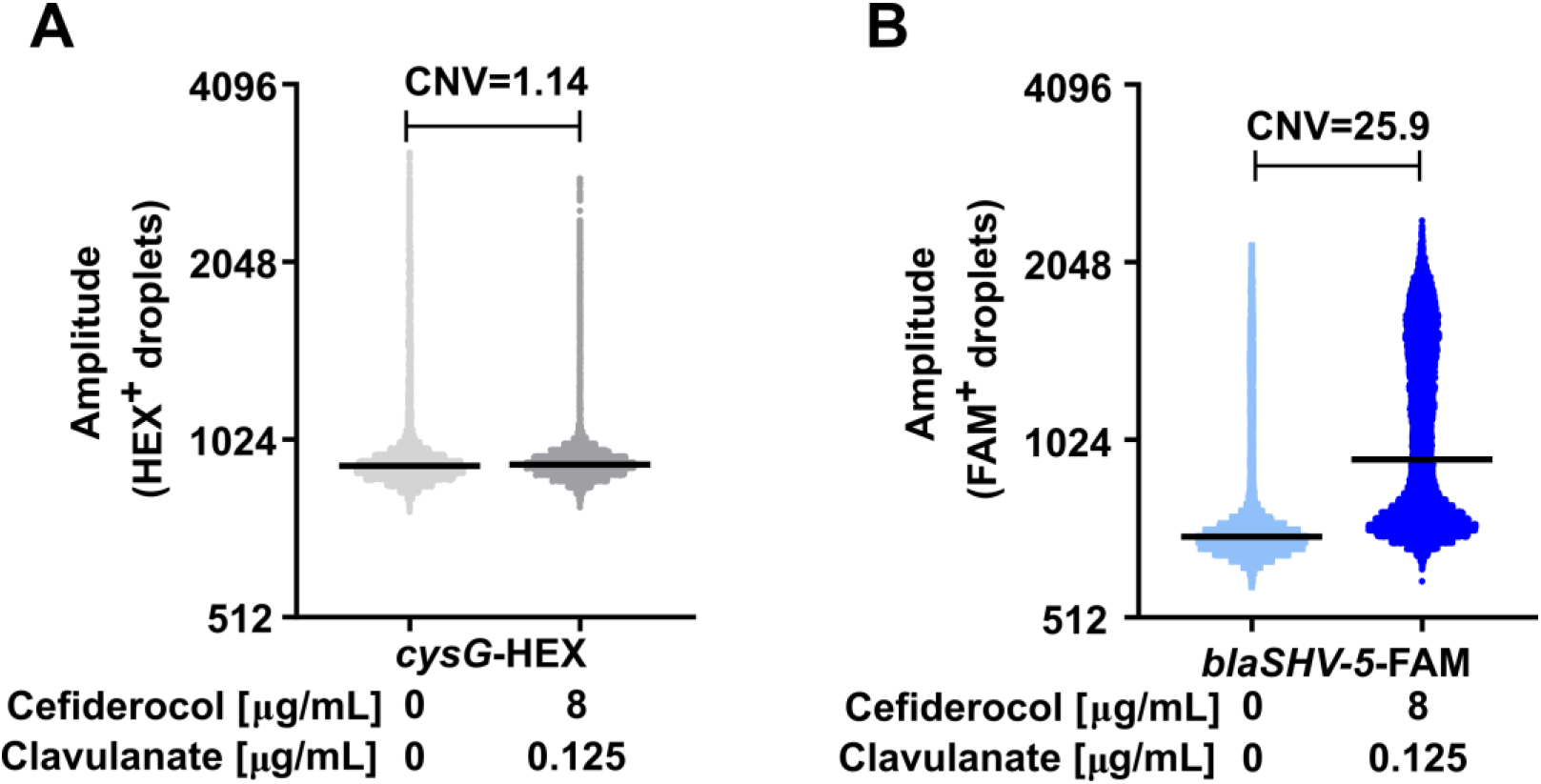
Droplet digital PCR analysis of *blaSHV-5* abundance confirms qPCR results. Droplet digital PCR was performed on gDNA isolated as part of the experiment in Figure 2E, from *Enterobacter* RS grown on MHA containing 0 or 8 µg/mL cefiderocol and 0.125 µg/mL clavulanate. Each DNA sample was partitioned into ∼20,000 droplets, PCR was performed, and amplification of the genes was detected for each droplet by quantifying the fluorescence of the HEX and FAM reporter probes. The probe internal to the PCR product amplifying (**A**) the housekeeping gene *cysG* was conjugated to the HEX fluorophore, and the (**B**) *blaSHV-5* probe was conjugated to the FAM fluorophore. Graphed are the raw amplitude values of each reporter from droplets containing both the HEX and FAM probes; >15,000 droplets per sample are shown. QuantaSoft Software uses a Poisson algorithm to quantify each gene as copies/µl input, and the copy number variation (CNV) is computed. The amplitude and CNV value for each pairwise comparison are shown above from a single replicate, representative of two biological replicates measured.

**Figure S7.**
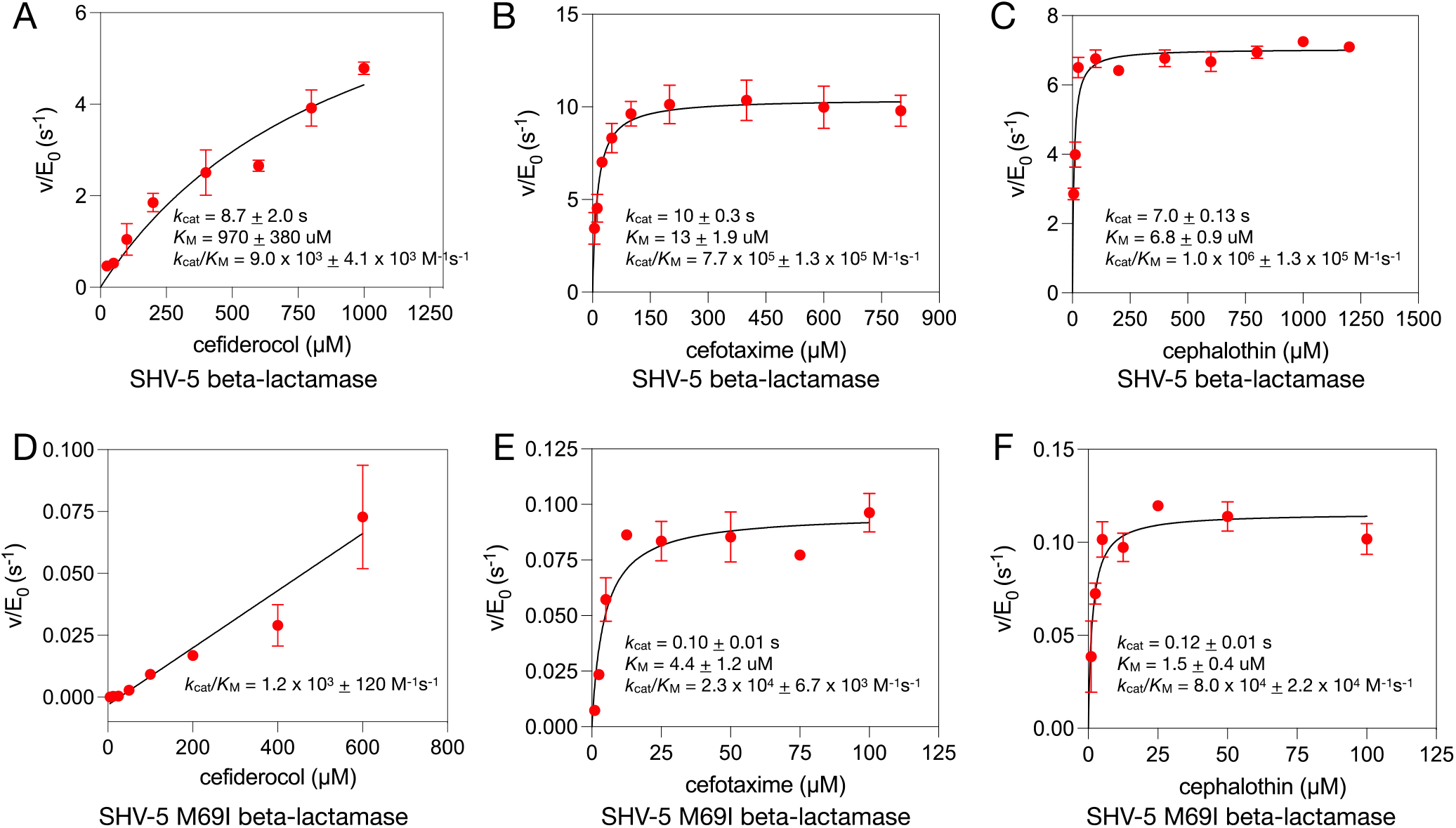
Michaelis-Menten steady-state kinetic parameter determinations for SHV-5 and SHV-5 M69I β-lactamases with cephalosporin substrates. (**A)** Enzyme initial velocity measurements of SHV-5 with increasing concentrations of cefiderocol. The data are fit to a hyperbola to yield *k*_cat_, *K*_M_, and *k*_cat_/K_M_ values, which are indicated. (**B)** Enzyme kinetic parameter determinations for SHV-5 and cefotaxime. (**C)** Enzyme kinetic parameter determinations for SHV-5 and cephalothin. (**D)** Enzyme kinetic parameter determination for SHV-5 M69I and cefiderocol. (**E**) Enzyme kinetic parameter determinations for SHV-5 M69I and cefotaxime. (**F**) Enzyme kinetic parameter determinations for SHV-5 M69I and cephalothin.

**Supplemental Figure 8.**
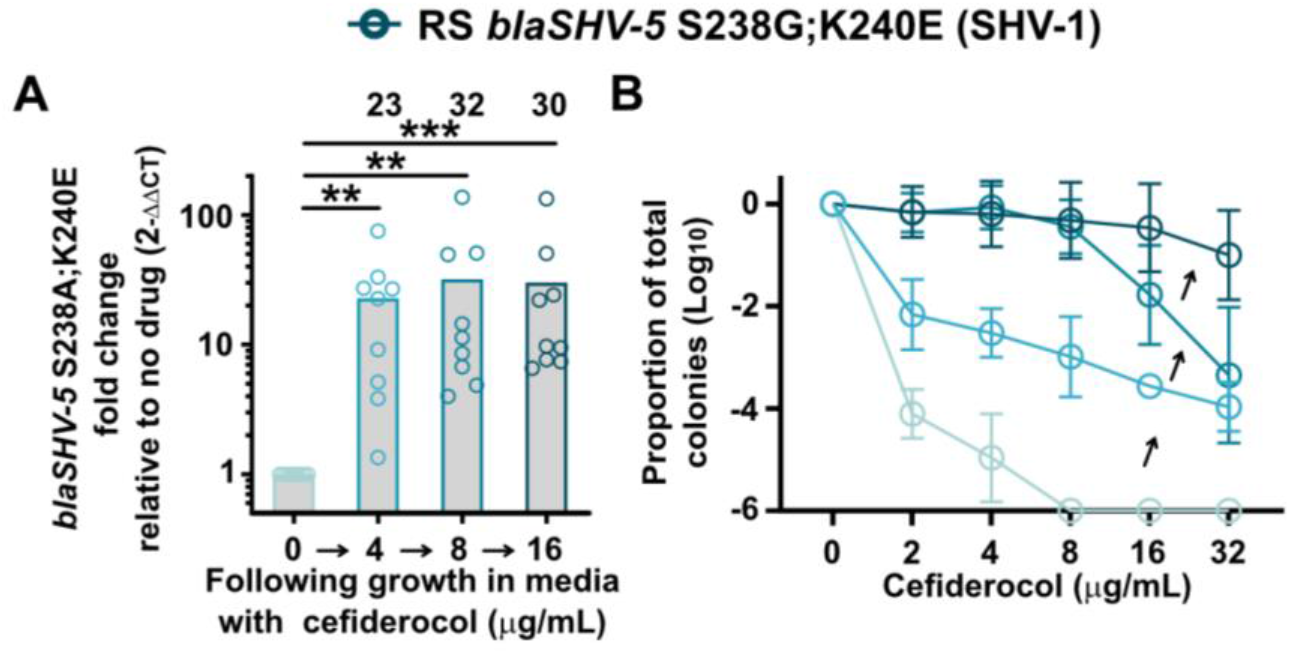
The magnitude of amplification of *blaSHV-1* is greater than *blaSHV-5* and capable of generating resistance. (**A**) Δ*blaSHV-5*::*blaSHV-5 S238G;K240E* (*blaSHV-5* S238G;K240E; which is SHV-1) abundance after resistant colonies were collected on MHA or MHA containing cefiderocol and grown in broth containing the same concentration indicated. (**B**) PAP of samples from panel (A). A-B show the mean from 3 independent experiments with 2-3 biological replicates each, with each dot indicating a biological replicate in A, and standard deviation shown in B. For A, ** indicates p<0.01,and *** p<0.001 by Kruskal-Wallis test with Dunnet’s correction for multiple comparisons.

### Beta-lactamase amplification in carbapenem-resistant *A. baumannii* (Figures S9-S11)

Having established that an ESBL causes cefiderocol HR in *Enterobacter* RS, we investigated whether beta-lactamase amplification might be a widespread mechanism to generate cefiderocol resistant subpopulations in HR isolates. As a first step, we attempted to determine which HR isolates might require a beta-lactamase to resist cefiderocol. If the resistant subpopulation in an HR isolate was absent when treated with cefiderocol in combination with a beta-lactamase inhibitor (BLI), then a beta-lactamase would be implicated. We performed PAP with cefiderocol alone or with the BLIs clavulanate or avibactam for each of the 27 carbapenem-resistant (CR) Enterobacterales and 65 CR-*Acinetobacter* from our surveillance collection that exhibited cefiderocol HR. Clavulanate had minimal effect for most isolates, consistent with its scope of inhibition being limited to only some members of class A and D beta-lactamases^1^. Avibactam, which inhibits class A, C, and some class D serine beta-lactamases, but not class B metallo-beta-lactamases, was effective at eliminating the resistant subpopulation in most isolates of each group, implicating serine beta-lactamases as being required for cefiderocol HR in these isolates (Figure S9).

In order to further investigate the role of beta-lactamases in cefiderocol HR, we selected representative isolates of CR-*Acinetobacter*, Mu1956 and Mu1984. We next assessed whether the resistant subpopulation in these strains was unstable by enriching it in cefiderocol and then passaging the strains into drug-free media. The resistant cells of each strain were enriched in cefiderocol and returned towards the baseline frequency subsequently in the absence of the drug, demonstrating the unstable HR phenotype in all three isolates (Figure S10A, D). To determine if the resistant subpopulation in each strain harbored beta-lactamase amplifications, we sequenced each isolate after growth with or without cefiderocol. When grown with cefiderocol, Mu1956 exhibited amplification of a 55.2 kbp region that included the *blaADC-30* gene, encoding an ESBL which is one of the most prevalent *Acinetobacter*-derived cephalosporinase (ADC, class C) variants^2^ (Figure S10B). The resistant subpopulation in Mu1956 was eliminated by addition of avibactam (Figure S10C). An amplicon with similar architecture including an ESBL was identified in Mu1984, with the *blaADC-33* gene encoding a related ADC^3^; this resistant subpopulation was also eliminated by avibactam (Figure S10E, F), consistent with the susceptibility of ADCs to avibactam^2^. Together, these data indicate that cefiderocol HR isolates can harbor resistant subpopulations with ESBL gene amplifications.

Data in Figure 2 suggest that with the concentration of cefiderocol held constant, cells required a dose-dependent increase in *blaSHV-5* to survive the increasing clavulanate concentration. We next tested whether this relationship is true with avibactam inhibition of the cefiderocol resistant subpopulation of *A. baumannii* Mu1956 and Mu1984. Each strain was grown with or without cefiderocol, as well as in media with cefiderocol and the highest concentration of avibactam in which the strain could survive. Growth in cefiderocol or cefiderocol/avibactam similarly enriched the cefiderocol resistant subpopulation (Figure S11A, C), but the average amplification of the *blaADC* genes was greater after growth containing avibactam (Figure S11B, D).

**Supplemental Figure 9.**
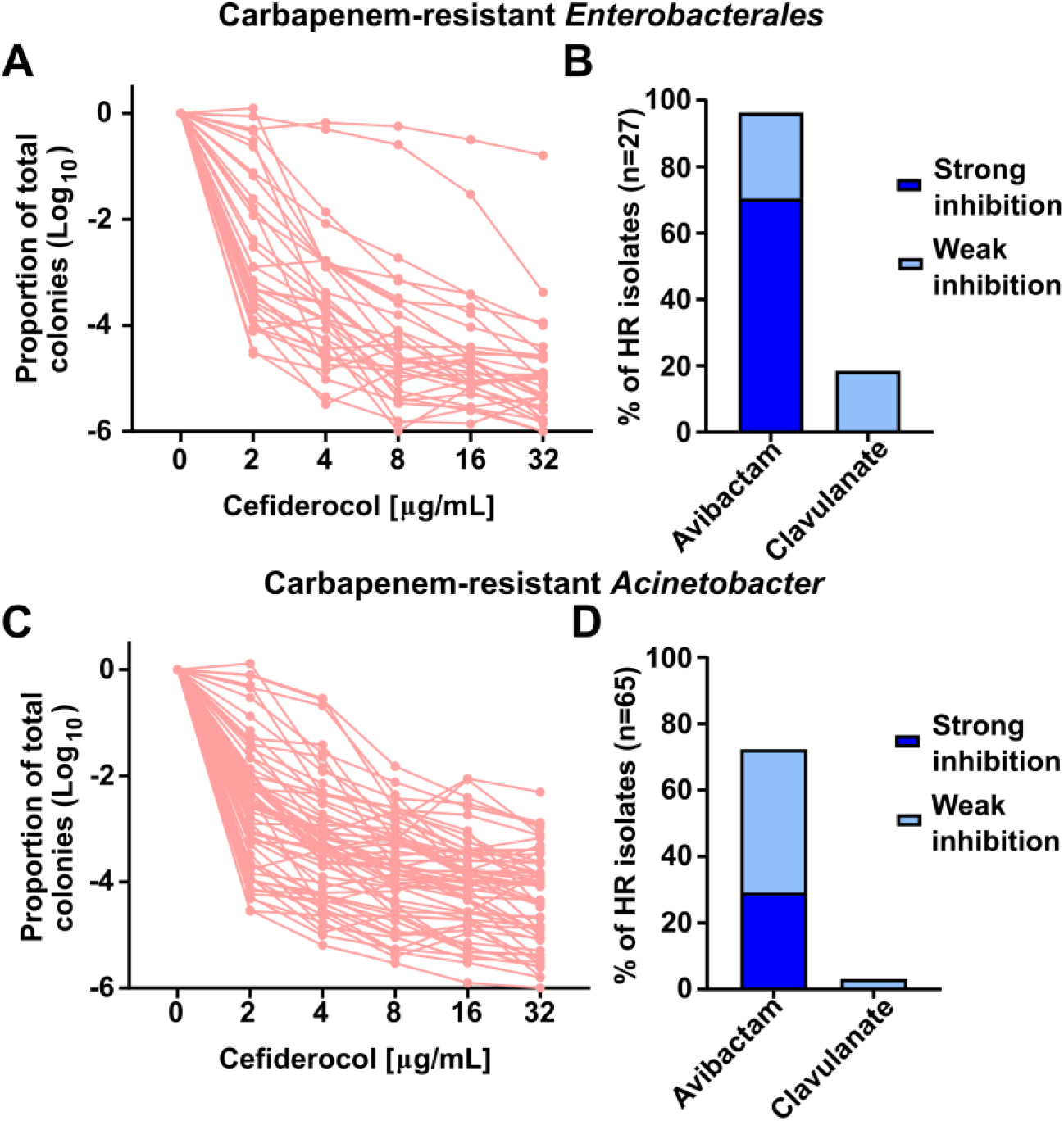
Cefiderocol heteroresistant CRE and CRAB clinical isolates are largely susceptible to avibactam inhibition. (**A**) Population analysis profile (PAP) of 27 carbapenem-resistant Enterobacterales (CRE) isolates identified as cefiderocol heteroresistant, where each line represents a unique isolate. (**B**) Results of PAP analysis for each isolate plated on cefiderocol plus avibactam or clavulanate. Inhibition was classified as “strong” if the presence of the respective beta-lactamase inhibitor caused the proportion of the surviving population to be below -6 log at ≤8 µg/mL cefiderocol, and was classified as “weak” if the presence of the inhibitor caused the proportion of the surviving population to be below -6 log at ≥16 µg/mL but not 8 µg/mL cefiderocol. The graphs are stacked bars. (**C-D**) Similar analyses as in **A-B** of 65 unique carbapenem-resistant *Acinetobacter baumannii* (CRAB) isolates.

**Supplemental Figure 10.**
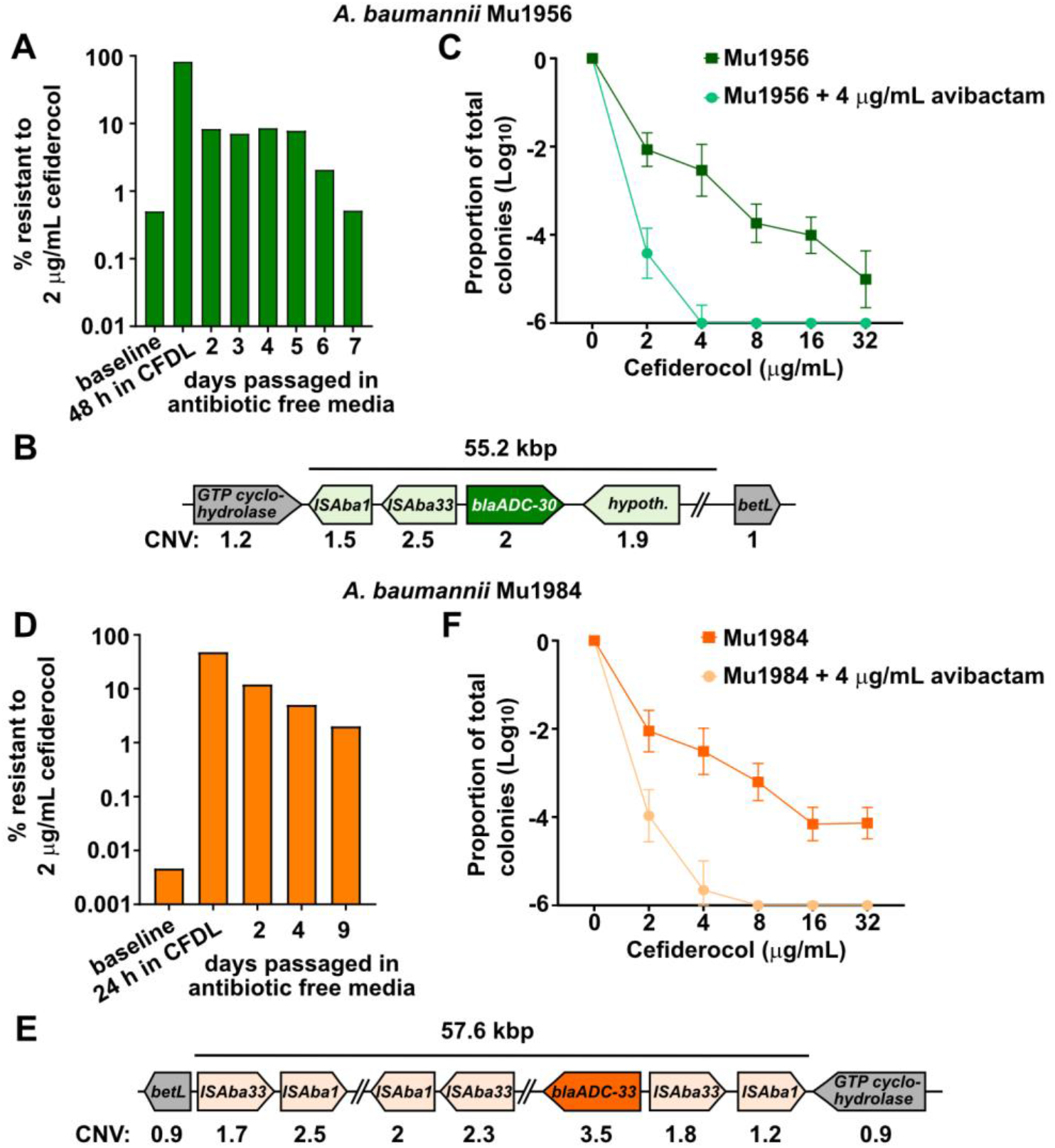
Cefiderocol resistant subpopulations of carbapenem-resistant *A. baumannii* contain amplified regions encoding beta-lactamases. (**A**) Quantification of the resistant subpopulations of *A. baumannii* Mu1956 in broth containing 32 µg/mL cefiderocol and subsequent passaging in antibiotic free media. (**B**) The region of the *A. baumannii* strain Mu1956 chromosome with amplification in the cefiderocol resistant population is shown, with copy number variation (CNV) shown below. (**C**) PAP of Mu1956 plated on cefiderocol or cefiderocol plus avibactam. (**D**) Quantification of the resistant subpopulations of *A. baumannii* Mu1984 in broth containing 32 µg/mL cefiderocol and subsequent passaging in antibiotic free media. (**E**) The region of the *A. baumannii* strain Mu1984 chromosome with amplification in the cefiderocol resistant population is shown, as is gene abundance. (**F**) PAP of Mu1984 plated on cefiderocol or cefiderocol plus avibactam. For **C** and **F**, the mean and standard deviation from two independent experiments with 3 or 6 biological replicates each is shown. In **B** and **E**, parallel lines indicate sequence regions omitted from the schematic.

**Supplemental Figure 11.**
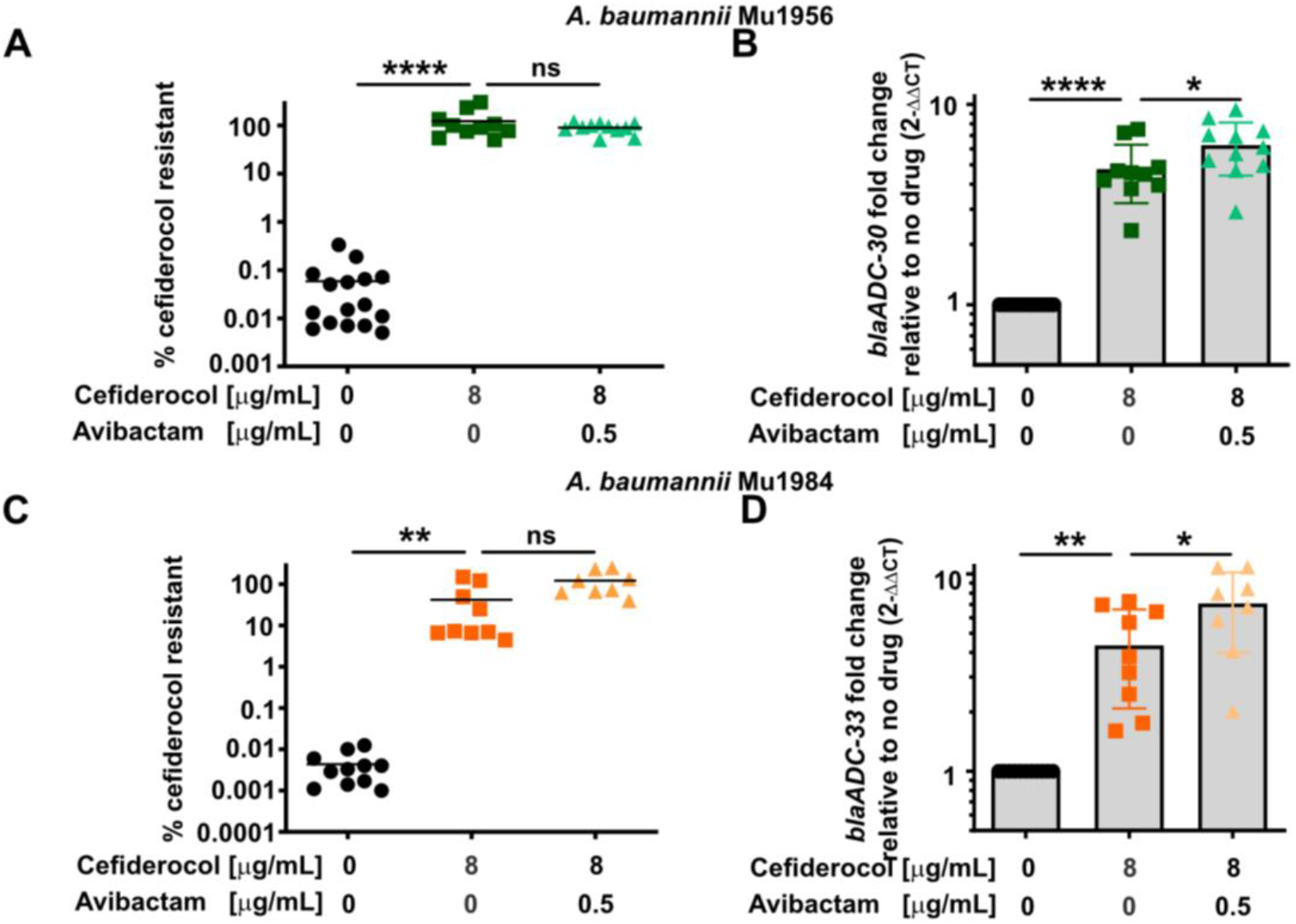
The magnitude of *A. baumannii* beta-lactamase amplification increases in the presence of a beta-lactamase inhibitor. (**A**) Quantification of Mu1956 survival on MHA containing 8 µg/mL cefiderocol after growth in media conditions indicated below the x axis. (**B**) Relative *blaADC-30* abundance normalized to *clpX* was quantified from the samples grown in broth in (**A**). (**C**) Quantification of Mu1984 survival on MHA containing 8 µg/mL cefiderocol after growth in media conditions indicated below the x axis. (**D**) Relative *blaADC-33* abundance normalized to *clpX* was quantified from the samples grown in broth in (**C**). Each symbol represents a biological replicate, with the mean indicated by a horizontal line. * indicates p<0.05, ** p<0.01, *** p<0.001, and ****p<0.0001 by one-way ANOVA [for A: F (2, 34) = 27.58, for B: F (2, 34) = 60.91, for C: F (2, 25) = 12.57 and for D: F (2, 25) = 20.30] with Sidak’s correction for multiple comparisons. (A-B) are from two independent experiments with 2-10 biological replicates each, and (C-D) from four independent experiments with 1-5 biological replicates each.

### ESBL copy number increase in response to greater antibiotic stress overcomes beta-lactamase inhibitors (Figure S12)

Mu1956 was HR to cefiderocol, but our screen found it susceptible to cefiderocol/avibactam (Figure S10C), suggesting the addition of avibactam could increase the utility of cefiderocol against this strain. However, based on our finding that beta-lactamase amplification can overcome the addition of an effective BLI (Figure 4), we investigated whether *blaADC-30* amplification could generate a population of cells resistant to the breakpoint concentrations of cefiderocol/avibactam. We first grew Mu1956 in broth containing 4 µg/mL cefiderocol plus 1 µg/mL avibactam, representing one quarter the breakpoint concentrations of each drug. Exposure to sub-breakpoint concentrations of drug is likely to occur for some cells in the population during infection. After a single exposure to sub-breakpoint cefiderocol/avibactam, *blaADC-30* gene frequency increased in most replicates (Figure S12A) and the surviving subpopulation became resistant to up to 16 µg/mL cefiderocol/4 µg/mL avibactam (Figure S12B) representing the breakpoint concentration of each drug^4^. These data highlight how rapidly an HR isolate can develop a subpopulation of cells resistant to the combination of cefiderocol plus an effective beta-lactamase inhibitor; one exposure to sub-inhibitory concentrations just under the breakpoint concentrations selected for a population with elevated *blaADC-30* amplification and enhanced resistance to the combination. These data highlight how beta-lactamase gene amplification threatens both current beta-lactam/BLIs as well as those in the development pipeline.

**Supplemental Figure 12.**
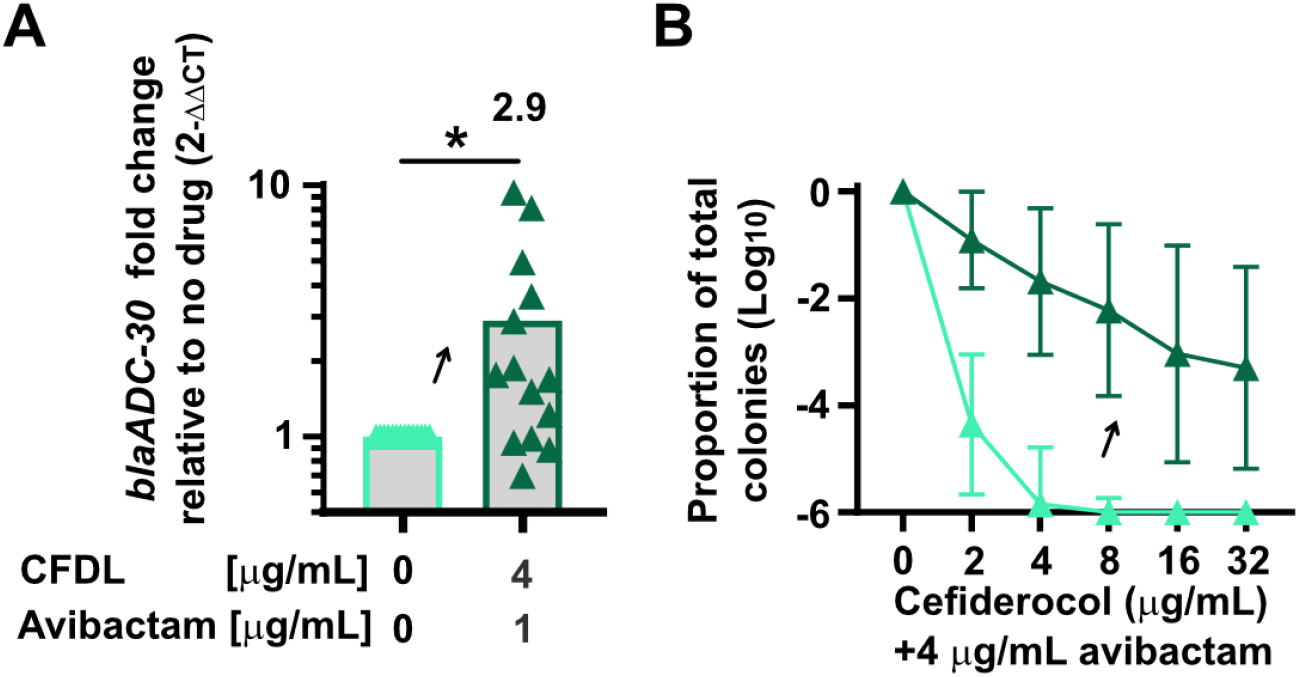
Cefiderocol/avibactam exposure increases amplification and results in a resistant subpopulation. (**A**) *blaADC30* abundance of *A. baumannii* Mu1956 after growth in broth containing cefiderocol and avibactam as indicated. (**B**) PAP of samples grown as indicated in **B.** Shown are results from three independent experiments with 3-6 biological replicates each with the mean shown. For **A**, each dot indicates a biological replicate and for B standard deviation is shown. For **A**, * indicates p<0.05 by paired two-tailed t-test, t=2.570, df=13.

**Supplemental Table 1.** Broth microdilution does not detect cefiderocol resistant subpopulations. The results of cefiderocol broth microdilution of samples grown in broth only (no cefiderocol) or in broth with cefiderocol. Shown are the results from at least 3 biological replicates each.

| Isolate | MIC (µg/mL)<br>No cefiderocol | MIC (µg/mL)<br>+ cefiderocol |
| --- | --- | --- |
| *Enterobacter* RS | 2 | >64 |
| *A. baumannii* Mu1956 | 4 | >64 |
| *A. baumannii* Mu1984 | 2 | >64 |

